# Revealing Acute Consequences of Rapid Protein Elimination at Individual Synapses using Auxin-Inducible Degron 2 Technology

**DOI:** 10.1101/2024.10.20.619267

**Authors:** Lilach Elbaum, Weixiang Yuan, Johannes P.-H. Seiler, Nadia Blom, Ya-Chien Chan, Ali Hyder Baig, Nils Brose, Simon Rumpel, Noam E. Ziv

## Abstract

A powerful approach to assess a protein of interest (POI) function is its specific elimination. Common knock-out and knock-down strategies, however, are protracted and often irreversible, challenging the assessment of acute or temporary consequences in the same cells and tissues. Here we describe the use of Auxin-Inducible Degron 2 (AID2) technology to study the real-time consequences of acute POI elimination in nerve cell synapses. We demonstrate its capacity in cultured neurons and *in vivo* to rapidly eliminate postsynaptic scaffold proteins fused at N-terminal, C-terminal, or nested sites to GFP derivatives or HaloTag. We show that acute PSD-95 or gephyrin elimination leads to the concomitant loss of AMPA or GABA_A_ receptors at the same synapses, and that, surprisingly, acute GKAP, but not PSD-95 elimination reduces postsynaptic scaffold size. Our findings highlight the utility of AID2 technology for rapidly eliminating synaptic POIs and studying real-time consequences in the same neurons and synapses.

## Introduction

A powerful approach to assess the functions and importance of a protein of interest (POI) in physiological and pathological processes is its specific elimination. For decades, constitutive or conditional knock-out approaches that act at the genome level have been considered the ‘gold standard’ approach in this regard. Most knock-out approaches, however suffer several inherent drawbacks. One is the irreversibility of the elimination step when carried out at the genomic level. The second is the protracted time course of POI elimination. Put differently, the time interval between the actual manipulation and its manifestation or assessment is typically long, due to the time course of organism development in the case of constitutive knock-outs or the long lifetimes of many proteins, particularly in neurons and synapses (Cohen and Ziv, 2019) in the case of conditional knock-outs. These protracted time courses leave ample time for slow ‘compensatory’ and adaptive processes that often mask the direct consequences of POI elimination and complicate interpretations. Furthermore, such protracted time scales pose a major challenge for experimental approaches aiming to follow the consequences of POI elimination in the same networks, cells, or sub-cellular compartments, such as synapses. To address these drawbacks and reduce the complexities associated with genetic knock-out approaches, alternative technologies focusing on the elimination of specific mRNAs (RNA interference, or ‘knock-down’) were developed (e.g. Hammond et al., 2001). But here again, POI elimination is typically slow, and incomplete in many instances. Moreover, off-target effects are not uncommon (Goel and Ploski, 2022).

Targeted protein degradation represents a categorically different approach to POI removal. Here, endogenous degradation systems, e.g. the ubiquitin-proteasome system or autophagic pathways, are harnessed to degrade a POI, often in an inducible fashion. Arguably, the best-known approaches include PROTACs (Proteolysis targeting chimeras) and molecular glue degraders (Tsai et al., 2024), small molecules that recognize a POI and recruit endogenous E3 ligases that subsequently target the POI for degradation by the ubiquitin-proteasome system. Unfortunately, devising such small molecules for specific targets is highly non-trivial, involving a combination of rational design and ‘brute force’ screening, with success not guaranteed.

An alternative way to achieve targeted protein degradation is to fuse the POI to a highly specific ‘degron’ - a short protein sequence that targets the POI for degradation upon induction (Natsume and Kanemaki, 2017; Bondeson et al, 2022). Although such approaches suffer from the need to express engineered POI variants, they are very powerful experimental tools, as evidenced by their increasing use, mainly in cell line settings. A promising system in this regard is the Auxin-Inducible Degron 2 (AID2; Yesbolatova et al., 2020). This system includes three components taken from a ubiquitous signaling cascade in plants, induced by plant Auxin hormones: (i) the *Arabidopsis*-derived mini-AID (mAID) degron, which is fused to the POI, (ii) a modified E3 ubiquitin ligase from rice (*Oryza sativa* transport inhibitor response 1, or OsTIR1), which forms a Skp1-Cul1-F-box (SCF) E3 ligase complex with endogenous components, and (iii) the Auxin derivative 5-phenyl-indole-3-acetic acid (5-Ph-IAA). In the presence of 5-Ph-IAA, the E3 complex binds to the mAID degron and ubiquitinates it, resulting in proteasomal degradation of the fusion protein. AID2 is a second-generation AID technology (Nishimura et al., 2009), based on mutation of OsTIR1 (F74G) and modification of its ligand, which resolved prior problems such as leaky degradation and dependence on high inducer concentrations.

Although potentially powerful, the use of AID2 technology in the field of neuroscience has been sparse (Yesbolatova et al., 2020; see also Nakano et al., 2019), and to the best of our knowledge, has not been used yet to study synapses. We therefore set out to examine the utility of AID2 technology for studying synapse biology, focusing on the effects of the acute elimination of core postsynaptic density (PSD) proteins on synapse properties. Specifically, we assessed in real-time, in the same cells and synapses, the consequences of acute elimination of the postsynaptic scaffold proteins PSD-95, GKAP (excitatory synapses), and Gephyrin (inhibitory synapses). We designed fusion proteins of PSD-95, GKAP and Gephyrin, each with the mAID degron and with GFP spectral variants or HaloTags to visualize them, and coexpressed these with OsTIR1 in cultured rat cortical neurons and in the mouse auditory cortex *in vivo*. We found that the mAID-tagged proteins fused to GFP derivatives or HaloTag proteins were eliminated rapidly, effectively, and reversibly upon induction with 5-Ph-IAA, when the degron was positioned C-terminally (PSD-95), N-terminally (Gephyrin, GKAP), or intramolecularly (Gephyrin). Importantly, even in the presence of endogenous wild-type variants of the same proteins, rapid elimination of degron-tagged proteins had discernable effects within the same time frames, including the loss of glutamate and GABA receptors from postsynaptic sites. Surprisingly, acute elimination of PSD-95 did not appear to affect PSD size, while elimination of GKAP caused significant PSD shrinkage. Our findings highlight the reliability and robustness of the mAID2 system for targeting synaptic POIs, and they demonstrate the utility of this system for functional studies on particular synaptic proteins *in-situ* by measuring real-time effects of their acute removal from individual synapses.

## Results

### PSD proteins are degraded rapidly by the AID2 system

We first validated published mAID2 tools that had previously been tested in cultured mouse hippocampal neurons (Yesbolatova et al., 2020). We obtained the original pAAV-hSyn-OsTIR1(F74G) plasmid from a public repository (see Methods), and moved the coding region from an AAV to a lentiviral backbone. This vector codes for OsTIR1(F74G) and, separated by a P2A sequence, EGFP fused to a C-terminal mAID degron (OsTIR1-P2A-EGFP:mAID; Fig S1A). When this construct is expressed in cultured rat cortical neurons, exposure to 200 nM 5-Ph-IAA is followed by the rapid (<2 h) disappearance of EGFP, in agreement with the original report (Yesbolatova et al., 2020; two separate experiments, 13 and 17 neurons, control and 5-Ph-IAA-treated, respectively; Fig. S1B,C).

To test the functionality of the AID2 system for degradation of synaptic proteins, we generated a lentiviral vector encoding a fusion protein of PSD-95, the fluorescent protein mTurquoise2 (Goedhart et al., 2010), and mAID (PSD-95:mTurq2:mAID; Fig. 1A). PSD-95 is a scaffold protein of the postsynaptic density of glutamatergic synapses which plays a crucial role in organizing synaptic components, in particular the confinement of multiple classes of glutamate receptors to postsynaptic membranes (reviewed in Won et al., 2017). When expressed in cultured rat cortical neurons, the PSD-95:mTurq2:mAID protein localized to postsynaptic sites along dendrites as previously observed for a PSD-95:mTurq2 variant lacking the mAID degron (Dvorkin and Ziv, 2016). To follow the 5-Ph-IAA induced degradation of PSD-95:mTurq2:mAID and the corresponding kinetics, cortical neurons were cotransduced with lentiviruses coding for PSD-95:mTurq2:mAID and lentiviruses coding for OsTIR1-P2A-EGFP:mAID, to introduce the essential OsTIR1(F74G) ubiquitin ligase and serve as positive controls. The preparations were mounted on a confocal microscope, connected to a slow perfusion system, and maintained at ∼37°C in an atmospheric environment of 5% CO_2_. After collecting baseline images for a few hours, 5-Ph-IAA (200 nM) was added to the preparations and the neurons were imaged at 1-h intervals. As shown for one neuron in Fig. 1B,C, and for 19 neurons from 3 independent experiments in Fig. 1D, exposure to 5-Ph-IAA was followed by the nearly complete elimination of PSD-95:mTurq2:mAID over 8-16 h, that followed an approximately exponential decay curve with a time constant of ∼3.7 h (see Methods).

**Figure 1.**
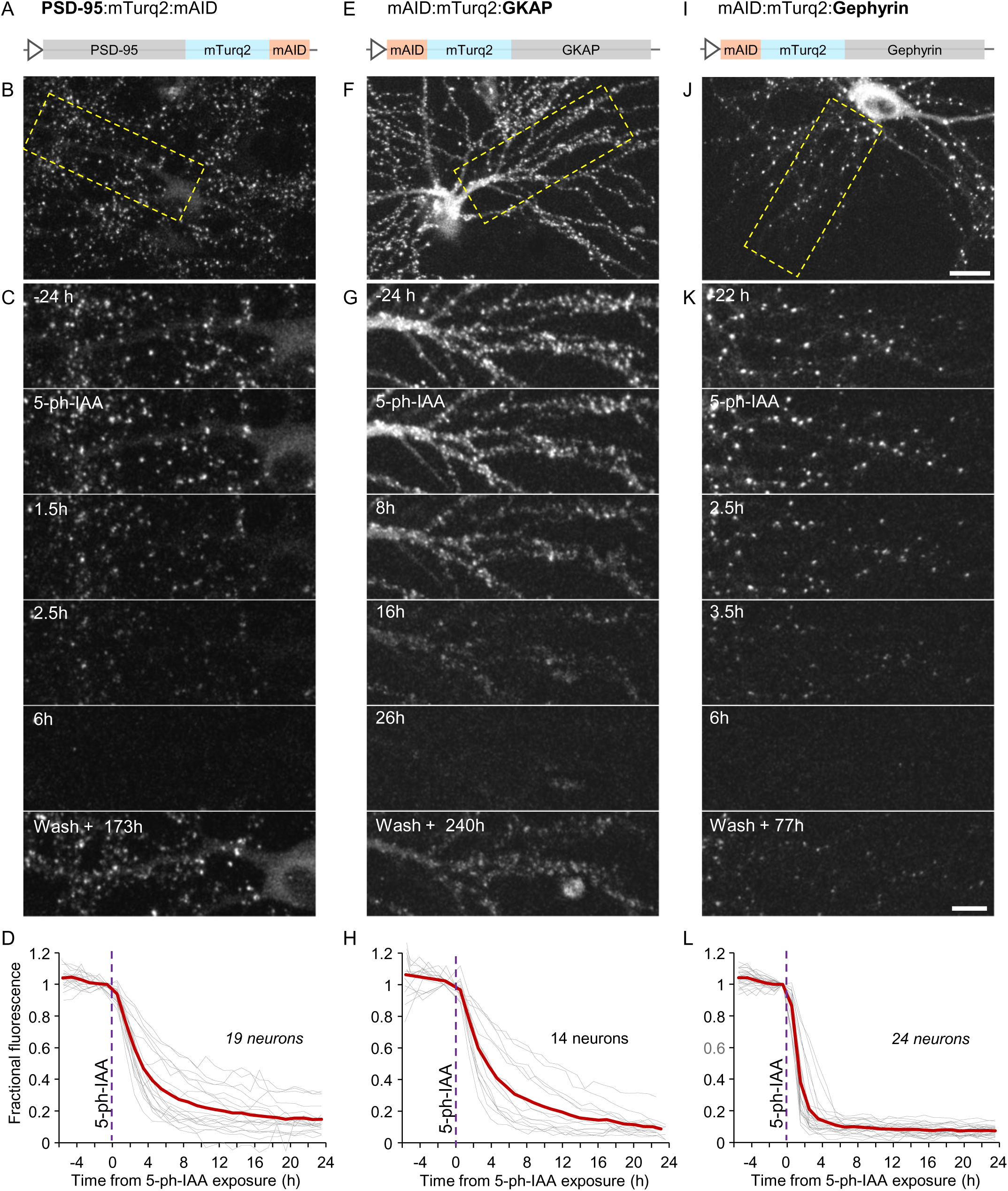
Rapid degradation of postsynaptic scaffold proteins fused to mTurquoise2 and a mAID degron. A) Illustration of the PSD:95, mTurquoise2 and mAID fusion protein (PSD-95:mTurq2:mAID). B) A rat cortical neuron in culture expressing PSD-95:mTurq2:mAID as well as OsTIR1-P2A-EGFP:mAID (see Supp. Fig. 1A; EGFP fluorescence not shown). C) Region in yellow rectangle in B at greater detail, before, after addition of 5-Ph-IAA, and after washing out the 5-Ph-IAA. Note the nearly complete loss of PSD-95:mTurq2:mAID fluorescence, and its recovery after several days. D) PSD-95:mTurq2:mAID fluorescence measured at 30 synapses of each neuron tracked throughout the experiments. Each thin gray line is the average fluorescence measured for the synapses of one neuron. Fluorescence values for each neuron were normalized to the fluorescence measured at the last time point before 5-Ph-IAA was added. Thick red line - population average (19 neurons from 3 independent experiments, 570 synapses in total). E-H) As in A-D for mAID:mTurq2:GKAP. 14 neurons from 2 independent experiments, 10-19 synapses per neuron, 199 in total). I-L) As in A-D for mAID:mTurq2:Gephyrin. 24 neurons from 3 independent experiments, 30 synapses per neuron, 720 synapses in total. Scale bars: 20 µm (B,F,J) and 10 µm C,G,K.

In the experiments described above, the mAID tag was fused to the C termini of EGFP and PSD-95:mTurq2. We also examined mAID2 efficacy when fused to the N-terminus of a POI (Yesbolatova et al., 2020). To that end, we created a fusion protein of the PSD protein GKAP (guanylate kinase-associated protein, also known as SAPAP1, DLGAP1 and DAP1) in which an N-terminally mTurquoise2-tagged rat GKAP was further N-terminally fused to mAID (mAID:mTurq2:GKAP; Fig. 1E). GKAPs are thought to bridge a membrane-proximal layer of scaffold proteins, including glutamate receptors and PSD-95, and a membrane distal layer, including SHANKs/ProSAPS and cytoskeletal linkers (Rasmussen et al., 2017). We chose GKAP because it was reported to be an important PSD organizer (Shin et al., 2012; Zhu et al., 2017; Zeng et al., 2018) and because N-terminal tagging with a fluorescent reporter protein was shown to maintain GKAP functionality (Kuriu et al., 2006). When mAID:mTurq2:GKAP was expressed in cortical neurons (together with OsTIR1-P2A-EGFP:mAID), mAID:mTurq2:GKAP assumed a postsynaptic localization (Fig. 1F,G), as previously seen with EGFP- and YFP-tagged GKAP (Yao et al., 2003; Kuriu et al., 2006). When followed as described above, the addition of 5-Ph-IAA led to the rapid elimination of mAID:mTurq2:GKAP (Fig. 1E-H) with a time constant similar to that of PSD-95:mTurq2:mAID (∼3.7 h; 14 neurons from two independent experiments).

PSD-95 and GKAP are well established components of excitatory glutamatergic synapses. Analogously, Gephyrin is considered to be the main organizer of the postsynaptic specialization at inhibitory GABAergic and glycinergic synapses, confining ionotropic GABA and Glycine receptors to postsynaptic sites (reviewed in Choii and Ko, 2015). To examine if mAID2 technology can be used to target scaffold proteins of GABAergic synapses, we tested a fusion protein of mAID, mTurquoise2 and Gephyrin, with the mAID and mTurquoise2 fused to the N-terminus of Gephyrin (mAID:mTurq2:Gephyrin; Fig. 1I), in accordance with prior studies (Calamai et al., 2009; Rubinski et al., 2015). When expressed in cortical neurons, mAID:mTurq2:Gephyrin assumed a punctate, dendritic expression pattern (Fig. 1J), identical to that observed for the same fusion protein lacking mAID (Rubinski et al., 2015). Again, in neurons coexpressing OsTIR1-P2A-EGFP:mAID, 5-Ph-IAA induced very rapid elimination of mAID:mTurq2:Gephyrin (Fig. 1 I-L) with an approximate time constant of 1.6 h (24 neurons from 3 separate experiments).

Finally, we examined whether mAID2 technology can also be used to rapidly degrade synaptic proteins labeled with a HaloTag (HT) (England et al., 2015), particularly when the HT is bound to fluorescent HT ligands, which could potentially interfere with proteasomal degradation. To that end we substituted mTurq2 with HT in PSD-95:mTurq2:mAID, obtaining PSD-95:HT:mAID (Fig. 2A). In this case, neurons expressing PSD-95:HT:mAID together with OsTIR1-P2A-EGFP:mAID were first exposed to the fluorescent ligand JF635-HT (Grimm et al., 2017), which led to the highlighting of postsynaptic sites (Fig. 2B). Exposure to 5-Ph-IAA led to the striking disappearance of ligand fluorescence, indicating that the fusion protein was degraded. The time constant of the elimination process (∼8.5 h; Fig. 2D), was somewhat longer than that observed for PSD-95:mTurq2:mAID (24 neurons from 3 separate experiments).

**Figure 2.**
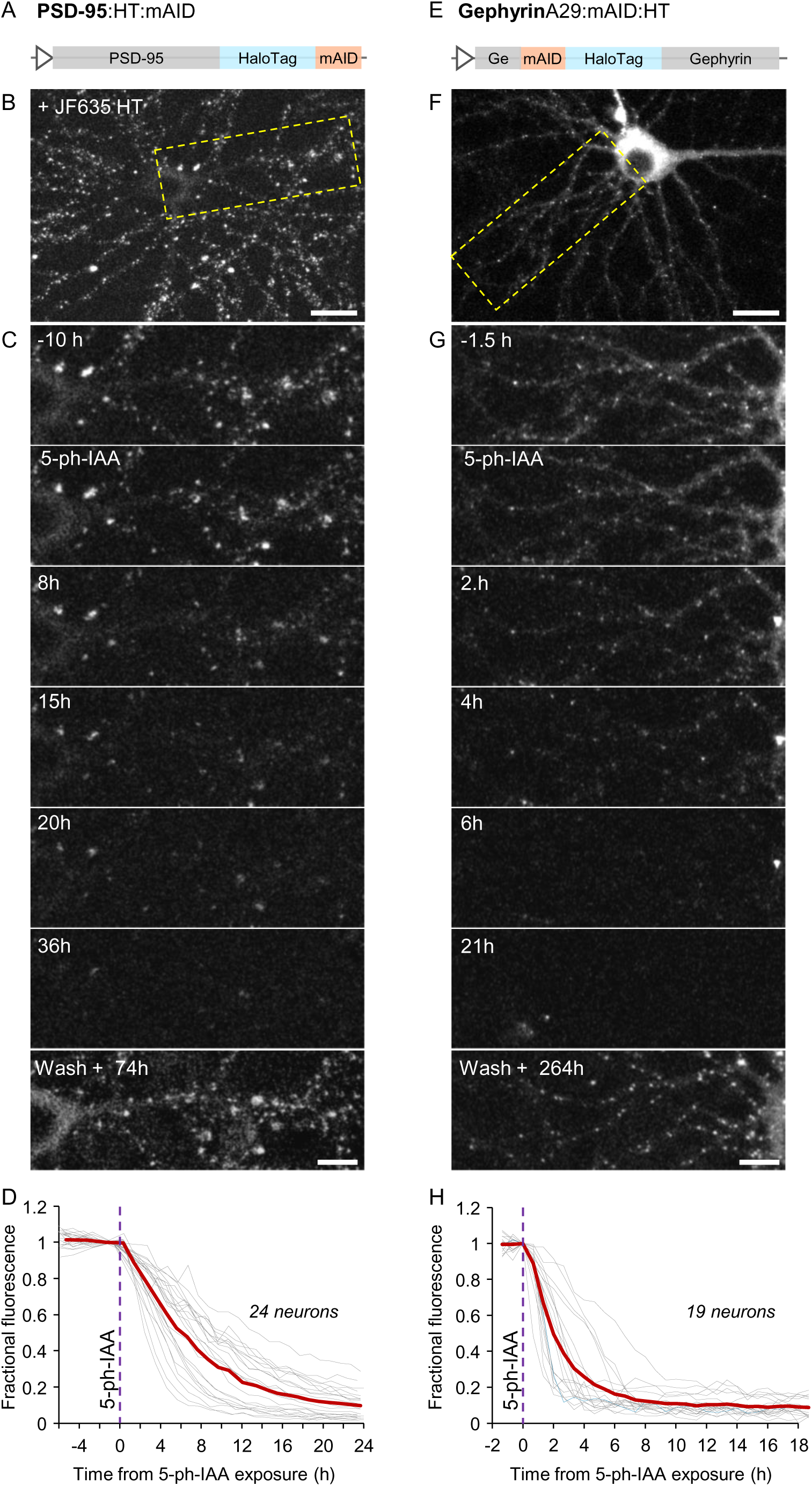
Rapid degradation of postsynaptic scaffold proteins fused to HaloTag protein and a mAID degron. A) Illustration of the PSD-95 - HaloTag - mAID fusion protein (PSD-95:HT:mAID). B) A rat cortical neuron in culture expressing PSD-95:HT:mAID as well as OsTIR1-P2A-EGFP:mAID (EGFP fluorescence not shown). PSD-95:HT:mAID was rendered visible by labeling with JF635-HT (100nM). C) Region in yellow rectangle in B at greater detail, before, after addition of 5-Ph-IAA, and after washing out 5-Ph-IAA. Note the nearly complete loss of JF635-HT fluorescence, and its recovery after several days (following a second labeling with JF635-HT). D) JF635-HT fluorescence measured at 11-30 synapses of each neuron (443 in total) tracked throughout the experiments. Each thin gray line is the average JF635-HT fluorescence measured for the synapses of one neuron. Fluorescence values for each neuron were normalized to the fluorescence measured at the last time point before 5-Ph-IAA was added. Thick red line is the population average (24 neurons from 3 independent experiments). E-H) As in A-D for GephyrinA29:mAID:HT. 18 neurons from 3 independent experiments. Scale bars: 20µm (B,F) and 10µm C,G.

Similarly, we generated a fusion protein of Gephyrin and a HaloTag, but decided to insert the mAID:HT sequence at a different site, after observing examples in which an antibody against gephyrin failed to recognize the N-terminal fusion protein variants described above (Supp. Fig. 2A). We therefore inserted the mAID:HT sequence between Asparagine 29 and Leucine 30 (GephyrinA29:mAID:HT; Fig. 2E), following the strategy Machado and colleagues used to create an mRFP-tagged Gephyrin knock-in mouse (Machado et al., 2011). As shown in Fig. 2F-H, GephyrinA29:mAID:HT assumed a dendritic, punctate expression pattern, similar to that observed with the original fusion proteins. Moreover, GephyrinA29:mAID:HT was now recognized by the anti-gephyrin antibody mentioned above (Supp. Fig. 2B). Exposure of neurons co-expressing GephyrinA29:mAID:HT and OsTIR1-P2A-EGFP:mAID caused the loss of ligand fluorescence, indicative of rapid degradation of the Gephyrin fusion protein (18 neurons from 3 separate experiments). Here too, elimination kinetics were somewhat slower that those observed for gephyrin fused to mTurq2 (time constant of ∼2.5 h).

As mentioned above, one of the main advantages of the mAID2 system is its reversibility. Indeed, when 5-Ph-IAA was washed out at the end of experiments, and the preparations were examined again after several days, clear recovery was observed for all the fusion proteins tested (Fig. 1C,G,K, Fig. 2C,G). Note that JF635-HT was re-added to the dish to visualize the HT fusion proteins. The time course of recovery was not followed systematically due to the slow turnover rates of synaptic proteins, and was sampled at single time points. Recovery, however, was very robust for all fusion proteins examined.

In summary, PSD proteins of excitatory and inhibitory synapses, labeled with either GFP variants or HT and N-terminally, C-terminally, or intramolecularly tagged with mAID are rapidly and reversibly degraded in the presence of OsTIR1(F74G) and 5-Ph-IAA.

### Dependence of elimination kinetics on expression levels

The variability of POI and OsTIR1 expression levels among neurons provided an opportunity to evaluate degradation rate dependence on mAID fusion protein and OSTIR1 expression levels. For these and subsequent experiments (see below) we created a new OsTIR1(F74G) expression vector (OsTIR1-P2A-mCherry) in which EGFP:mAID was replaced with mCherry (without the mAID degron, but still separated from OsTIR1 by a P2A sequence) which was coexpressed with PSD-95:mTurq2:mAID and exposed to 5-Ph-IAA as described above. We then measured, on a neuron by neuron basis, the initial rate of synaptic PSD-95:mTurq2:mAID degradation as a function of (i) initial PSD-95:mTurq2:mAID fluorescence, and (ii) mCherry fluorescence, the latter serving as a proxy for OsTIR1 expression levels. Initial rates of synaptic PSD-95:mTurq2:mAID degradation were estimated as linear fits to PSD-95:mTurq2:mAID fluorescence during the first 5 time points following 5-Ph-IAA addition. As shown in Supp. Fig. 3A, PSD-95:mTurq2:mAID degradation rates (in absolute fluorescence units per h) correlated well with PSD-95:mTurq2:mAID expression levels (r = 0.75), as might be expected, and also with OsTIR1 expression levels (r = 0.65; Supp. Fig. 3B). This sensitivity to OsTIR1 expression levels probably explains some of the variability in the degradation curves of individual neurons. It also indicates that OsTIR1 expression level, rather than those of the endogenous proteins needed to form active SCF E3 ligase complexes, is a limiting factor that dictates POI degradation rates.

### Degradation of mAID fusion proteins *in-vivo*

In their original study, Yesbolatova and colleagues (2020) demonstrated the utility of the AID2 system to drive mAID-EGFP degradation in various organs of transgenic mice following intraperitoneal 5-Ph-IAA injection. mAID-EGFP degradation was also observed in whole brains, albeit to a somewhat lesser extent (∼50%) than that observed in other organs (see also Makino-Itou et al., 2024). To examine the *in-vivo* efficacy of this system at the single neuron level, we first packaged the OsTIR1-P2A-EGFP:mAID vector from the original study into an AAV capsid (see Methods) and then stereotaxically injected it into the auditory cortex of wildtype mice together with a second AAV vector, coding for Histone 2B fused to mCherry (H2B:mCherry). This combination led to the expression of a cytosolically expressed mAID:EGFP, together with a stably expressed nuclear reference marker (Fig. 3A,B). After a recovery period, we implanted the injected mice with a cranial window and conducted longitudinal two-photon imaging, which allowed a detailed assessment of the time course of the degradation and recovery Specifically, we imaged immediately before (-1, -0.5 h) and repeatedly after (1,3,6,9,24,48 and 72 h) intraperitoneal administration of 10mg/kg 5-Ph-IAA (5-Ph-IAA group; 3 mice) or saline (sham group; 2 mice). In line with the original study reporting a decrease of GFP fluorescence by ∼50% after sacrificing mice 6 hours after injection (Yesbolatova et al., 2020), we observed a substantial decrease of cytosolic mAID:EGFP fluorescence after injection of 5-Ph-IAA, but not with saline (Fig. 3B; 91 and 111 neurons, respectively). In our experiments, the loss of mAID:EGFP fluorescence in cell bodies and the neuropil became apparent already within one hour and reached a maximal reduction of approximately 90% after 6 h. Moreover, following this initial reduction, a recovery of fluorescence was observed over the course of about three days (Fig. 3C). These observations demonstrate the efficient yet reversable depletion of a cytosolic protein using the AID2 system in individual neurons *in-vivo*.

**Figure 3.**
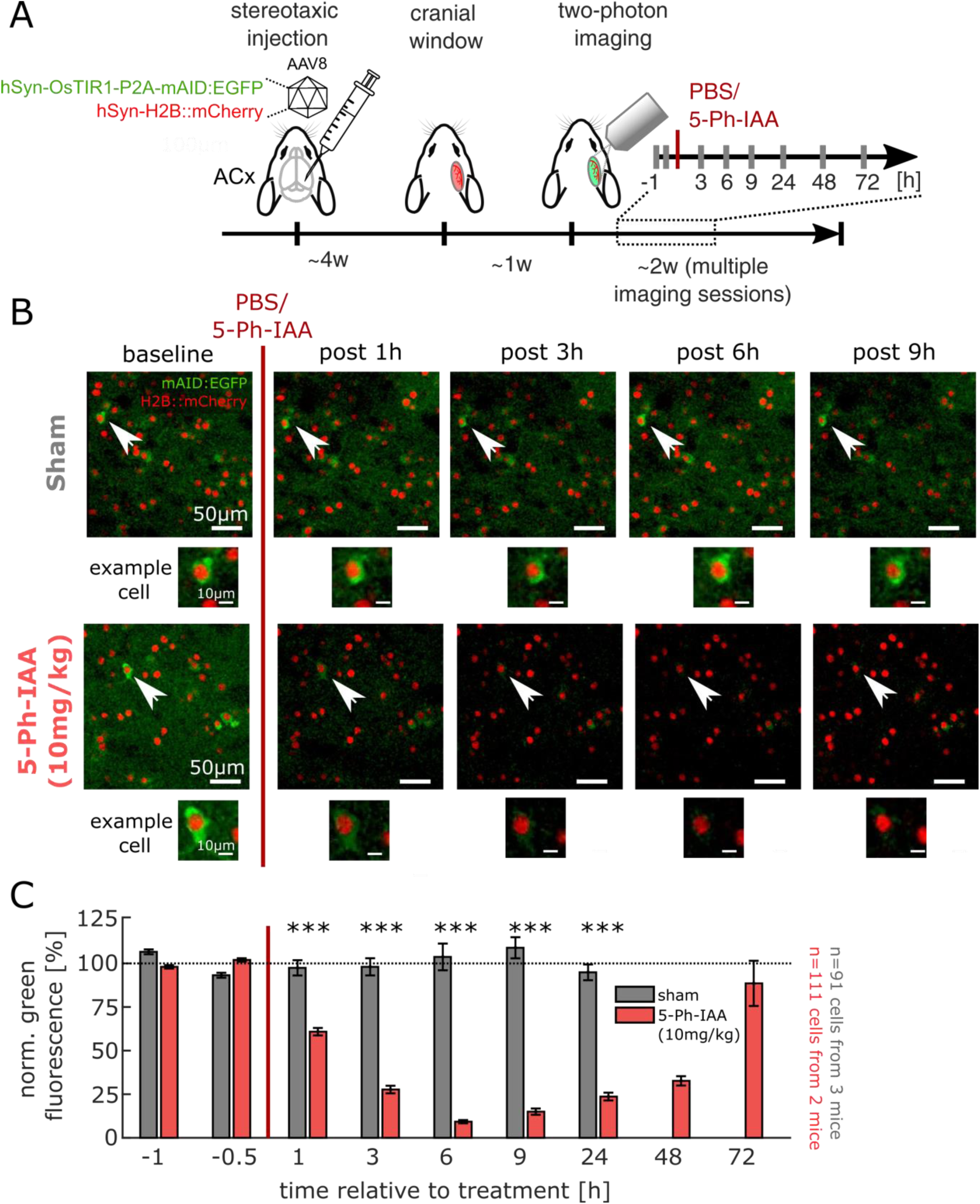
AID2-mediated elimination and recovery of a soluble protein *in vivo*. **A**) Flow of experiment. C57BL/6J mice (n = 3) were injected with AAV vectors to cytosolically express mAID:GFP, OsTir1 and a stable nuclear reference marker (H2B:mCherry) in the auditory cortex. Animals underwent cranial window implantation and longitudinal two-photon imaging after sham (PBS) or 5-Ph-IAA treatment. **B**) Exemplary fields of view, showing stable mAID:EGFP fluorescence after sham treatment and vastly depleted mAID:EGFP fluorescence after auxin treatment. **C**) Quantification of the normalized cytosolic mAID:EGFP fluorescence (see Methods) over the course of imaging (n=91 sham neurons vs. 111 5-Ph-IAA treated neurons), showing a swift depletion over the course of few hours and a subsequent recovery after approximately three days (***: p<0.001, Wilcoxon rank sum test of sham vs. 5-Ph-IAA).

We then set out to explore the utility of the AID2 system to degrade a synaptic protein *in-vivo*. Specifically, we injected wild type mice with a mixture of three AAV vectors in the auditory cortex: (i) hSyn-PSD95:HT:mAID to express a degradable PSD-95 variant, similar to that used for the cell culture experiments of Fig. 2A-D; (ii) hSyn-OsTIR1-P2A-NLS:tagBFP2 to allow the OsTIR1-dependent depletion of the mAID-tagged PSD-95, and (iii) hSyn-PSD95.FingR:EGFP-CCR5TC to label the exogenous as well as the endogenous PSD-95 (Gross et al., 2013). This set of vectors allowed FingR-mediated labeling of overall PSD-95 and a specific HT-mediated labeling of the exogenous, potentially degradable PSD-95 variant. After a period of three to four weeks to reach stable expression labels, mice were either intraperitoneally injected with saline (sham group, n=3) or with 10mg/kg 5-Ph-IAA (5-Ph-IAA group, n=4). Six hours after treatment, animals were sacrificed, their brains were extracted and fixed. Subsequently, brain slices were prepared, stained with the HaloTag ligand JF635-HT (Grimm et al., 2017) and mounted for confocal imaging (Fig. 4A; see Methods). As shown in Fig. 4B, all fluorescent markers showed robust expression. As expected, in slices from the sham group, PSD-95-associated FingR and HaloTag fluorophores exhibited punctate expression patterns with substantial overlap, likely corresponding to excitatory synapses. Importantly, in slices from the 5-Ph-IAA treated animals, JF635-HT fluorescence was significantly reduced to approximately 60% of the signal intensity of the fluorescence range between sham treated and non JF635-HT stained control slices (Fig. 4B,C). This drop in fluorescence concurs with the ∼50% reduction in PSD-95:HT:mAID observed in culture after 6 h (Fig. 2D). In contrast, treatments with 5-Ph-IAA only modestly affected the PSD-95 FingR signal (Fig. 4D), indicating that endogenous PSD-95 levels were largely unaltered.

**Figure 4.**
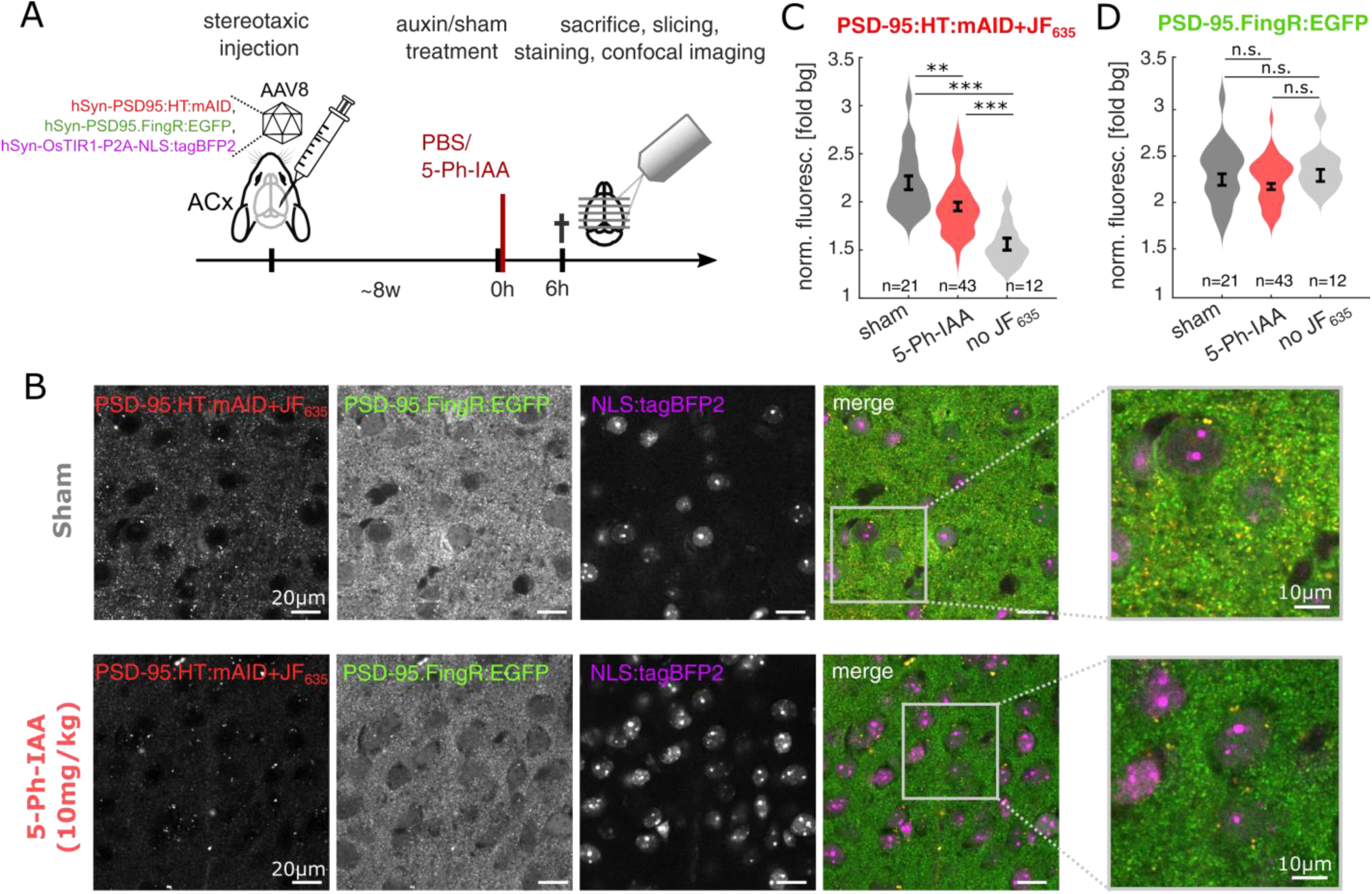
AID2-mediated elimination and recovery of a synaptic scaffold protein *in vivo***. A**) Flow of experiment. After stereotaxic injection and stable expression of the exogenous proteins, mice were treated with (sham) or 5-Ph-IAA (auxin). After 6 h, mice were sacrificed and their brains were extracted, fixed and stained with the HaloTag ligand JF635-HT before starting confocal imaging (n=3 sham mice vs. 4 5-Ph-IAA mice). **B**) Exemplary fields of view, showing all three imaging channels in a sham and 5-Ph-IAA treated animal. Right: Magnified image, showing the punctate pattern of PSD-95 signals, expected to correspond to excitatory synapses. The PSD-95:HT:mAID signal is decreased after the 5-Ph-IAA treatment compared to the sham treatment. **C**) Quantification of the normalized PSD-95:HT:mAID fluorescence in the neuropil, showing a significant signal depletion after 5-Ph-IAA injection (n indicates the number of FOV per condition; **: p<0.01, ***: p<0.001 in Wilcoxon rank sum test). Note that degradation of PSD-95:HT:mAID may not have reached its maximum at this six hours observation period (see Fig. 2D). **D**) Equivalent quantification for the PSD-95:FingR:EGFP signal, showing a slight, but non-significant drop in intensity after 5-Ph-IAA treatment.

These experiments thus confirmed the effective depletion of a cytosolic protein and a synaptic protein in neurons *in vivo* using AID2 technology.

### Acute synaptic scaffold protein elimination leads to receptor loss at the same synapses

PSD-95 has been shown to play crucial direct and indirect roles in anchoring glutamate receptors, and in particular AMPA type ionotropic receptors (AMPARs) at postsynaptic membranes (reviewed in Bessa-Neto and Choquet, 2023). We therefore assessed the impact of PSD-95 elimination on AMPAR content at the same synapses. To visualize AMPARs we expressed a fusion protein of GluA2 and Super Ecliptic pHluorin (e.g. Ashby et al., 2004; Kopec et al., 2006) previously used in our hands (SEpH:GluA2; Zeidan et al., 2012). Such fusion proteins effectively report outward facing AMPARs located in the neuronal membrane, as the fluorescence of receptors within typically acidic intracellular organelles is quenched. For these experiments, we used OsTIR1-P2A-mCherry instead of OsTIR1-P2A-EGFP:mAID to avoid spectral overlap with Super Ecliptic pHluorin. We then triple expressed PSD-95:mTurq2:mAID, SEpH:GluA2 and OsTIR1-P2A-mCherry in cortical neurons in primary culture, and followed individual postsynaptic sites as described above, before and after addition of 5-Ph-IAA. As shown in Fig. 5A, PSD-95:mTurq2:mAID and SEpH:GluA2 puncta exhibited excellent colocalization, with comparisons of PSD-95:mTurq2:mAID and SEpH:GluA2 on a synapse to synapse basis revealing a high correlation between the two (r= 0.64; p = 7.6*10^-55^; 455 synapses from 18 neurons from 3 experiments). The addition of 5-Ph-IAA and the consequential loss of PSD-95:mTurq2:mAID was associated with a ∼25% loss of SEpH:GluA2 fluorescence on average (Fig. 5C,E), with loss kinetics closely following those of PSD-95:mTurq2:mAID. As SEpH:GluA2 tends to photobleach, we also measured SEpH:GluA2 (and PSD-95:mTurq2:mAID) in some neurons only once every 12 h; the degree of SEpH:GluA2 fluorescence loss, however, was nearly identical (Fig. 5B-D). Moreover, comparison with neurons in the same experiments that expressed SEpH:GluA2 but not PSD-95:mTurq2:mAID (Fig. 5D) confirmed that the loss of SEpH:GluA2 fluorescence associated with PSD-95:mTurq2:mAID degradation was statistically significant (p=1.8*10^-4^; 18 and 10 neurons, respectively). Thus, even in the presence of endogenous PSD-95 (see below), rapid elimination of exogenous PSD-95 led to the rapid loss of AMPARs at the same synapses, presumably due to receptor diffusion away from synaptic sites and possibly receptor endocytosis (Bessa-Neto and Choquet, 2023).

**Figure 5.**
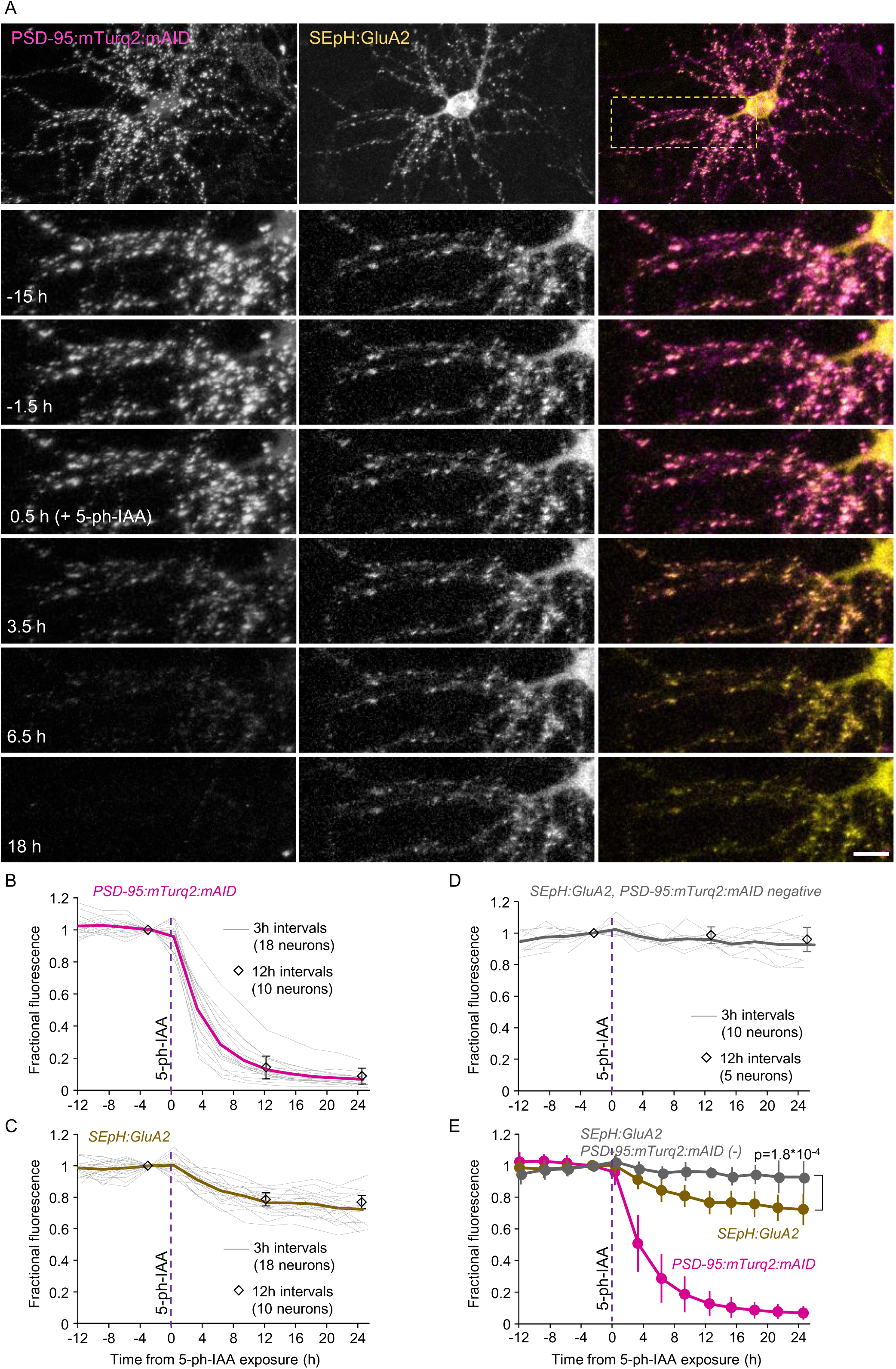
Acute elimination of PSD-95:mTurq2:mAID is followed by loss of AMPARs at the same synapses. **A)** Top panels: A rat cortical neuron in culture co-expressing PSD-95:mTurq2:mAID, SEpH:GluA2 and OsTIR1-P2A-mCherry (not shown). Bottom panels: Region in yellow rectangle at greater detail, before, and after addition of 5-Ph-IAA. Note that the near complete loss of PSD-95:mTurq2:mAID is associated with a modest reduction in SEpH:GluA2 fluorescence. Scale bars: 10 µm. **B)** PSD-95:mTurq2:mAID fluorescence measured at 16-41 synapses of each neuron (455 in total) tracked throughout the experiments. Each thin gray line is the average fluorescence measured for the synapses of one neuron (18 neurons from 3 experiments). Fluorescence was normalized to fluorescence measured at time point just before 5-Ph-IAA addition. Thick magenta line is the population average. A subset of neurons was imaged only once every 12 (instead of 3) h to minimize potential confounds related to photobleaching (open diamonds; 10 neurons from the same 3 experiments). **C)** changes in SEpH:GluA2 fluorescence at the same synapses and neurons of B. Thick brown line is the population average. **D)** SEpH:GluA2 fluorescence measured at 14-30 synapses of neurons positive for SEpH:GluA2 and OsTIR1-P2A-mCherry but negative for PSD-95:mTurq2:mAID (222 in total). Each thin gray line is the average fluorescence measured for the synapses of one neuron (10 neurons from 3 experiments). Thick gray line is the population average Open diamonds represent measurements made in a subset of cells imaged only once every 12 h (5 neurons from the same experiments). **E)** Pooled data. All error bars are standard deviations, not SEM. Test for difference between PSD-95:mTurq2:mAID positive and negative cells – unpaired t-test, without assuming equal variances; applied to data obtained at last time point.

It should be noted that even after the nearly complete loss of PSD-95:mTurq2:mAID from postsynaptic sites, substantial AMPAR confinement was still observed at the same sites (Fig. 5A,C,E). We therefore measured how substantial the lost PSD-95:mTurq2:mAID pool was, compared to presumably unaffected endogenous PSD-95 pools. To that end, we compared synaptic PSD-95 levels in PSD-95:mTurq2:mAID expressing and naïve neurons using quantitative immunocytochemistry. We found that PSD-95 levels at synapses expressing PSD-95:mTurq2:mAID were approximately 3-fold higher than those at synapses without PSD-95:mTurq2:mAID (24 and 31 fields of view from 2 separate experiments; Supp. Fig. 4). This substantial overexpression, a result of the high viral titers needed to successfully triple infect individual neurons, indicated that 5-Ph-IAA induced the loss of at least 75% of synaptic PSD-95.

In comparison to the significant loss of synaptic PSD-95, AMPAR loss was comparatively modest. This could be interpreted in several fashions. At one extreme, all synapses might have lost a similar and modest fraction of their receptor contents. At the other, the average value reported above might mask a broad range of receptor losses that differed from one synapse to another. Indeed, prior knockout and knockdown experiments indicate that AMPAR loss occurs at individual synapses in all-or-none fashion rather than as a uniform loss of some AMPARs from every synapse (Levy et al., 2015). To address this question, we compared, on a synapse by synapse basis, the loss of SEpH:GluA2 fluorescence to the loss of PSD-95:mTurq2:mAID fluorescence after a 15 h 5-Ph-IAA exposure period, using non-normalized fluorescence values, as these expected to scale linearly with fusion protein quantity. At the population level, SEpH:GluA2 fluorescence loss was positively correlated with PSD-95:mTurq2:mAID loss (r= 0.43, 455 synapses from 18 neurons from 3 experiments; Supp. Fig. 5A). A similar result was obtained when the correlation was calculated separately for each neuron (0.39 ± 0.22; mean ± standard deviation, respectively). Yet at the individual synapse level, SEpH:GluA2 loss was quite variable, with a considerable number of synapses showing no loss and even some gain of SEpH:GluA2 fluorescence (Supp. Fig. 5A, D). Comparisons with PSD-95:mTurq2:mAID fluorescence signals of similar magnitude indicate that this variability is not merely measurement noise (see Supp. Fig. 5A-C for further details). Although we observed no overt bimodality in the fraction of receptor loss (Supp. Fig. 5D), these findings argue against uniform receptor loss and indicate that synaptic AMPAR contents are only partially dictated by PSD-95 contents at individual synapses (see Discussion).

As mentioned above, Gephyrin, considered the main organizer of the GABAergic postsynaptic specialization, confines GABA receptors to postsynaptic sites. To examine if the rapid degradation of Gephyrin is also associated with receptor loss at the same synapses, we triple expressed, in cortical neurons, GephyrinA29:mAID:HT, OsTIR1-P2A-mCherry, and a fusion protein of GABA_A_ Receptor subunit α_2_ and Super Ecliptic pHluorin (SEpH:GABA_A_Rα_2_) previously shown to localize well to GABAergic synapses in cultured neurons and *in vivo* (Tretter et al., 2008; Nakamura et al., 2016). As shown in Fig. 6A, GephyrinA29:mAID:HT and SEpH:GABA_A_Rα_2_ colocalized extremely well at individual synapses. Quantitatively, comparing GephyrinA29:mAID:HT and SEpH:GABA_A_Rα_2_ on a synapse to synapse basis revealed a high correlation between the two (r= 0.64; p = 1.75*10^-105^; 900 synapses from 18 neurons from 3 experiments). We then followed individual synapses as described above, before and after addition of 5-Ph-IAA. The consequential loss of GephyrinA29:mAID:HT was associated with a ∼20% reduction of SEpH:GABA_A_Rα_2_ fluorescence (Fig. 6A-D). Here too, comparison with neurons that did not express GephyrinA29:mAID:HT, revealed that receptor loss was statistically significant (p = 0.003).

**Figure 6.**
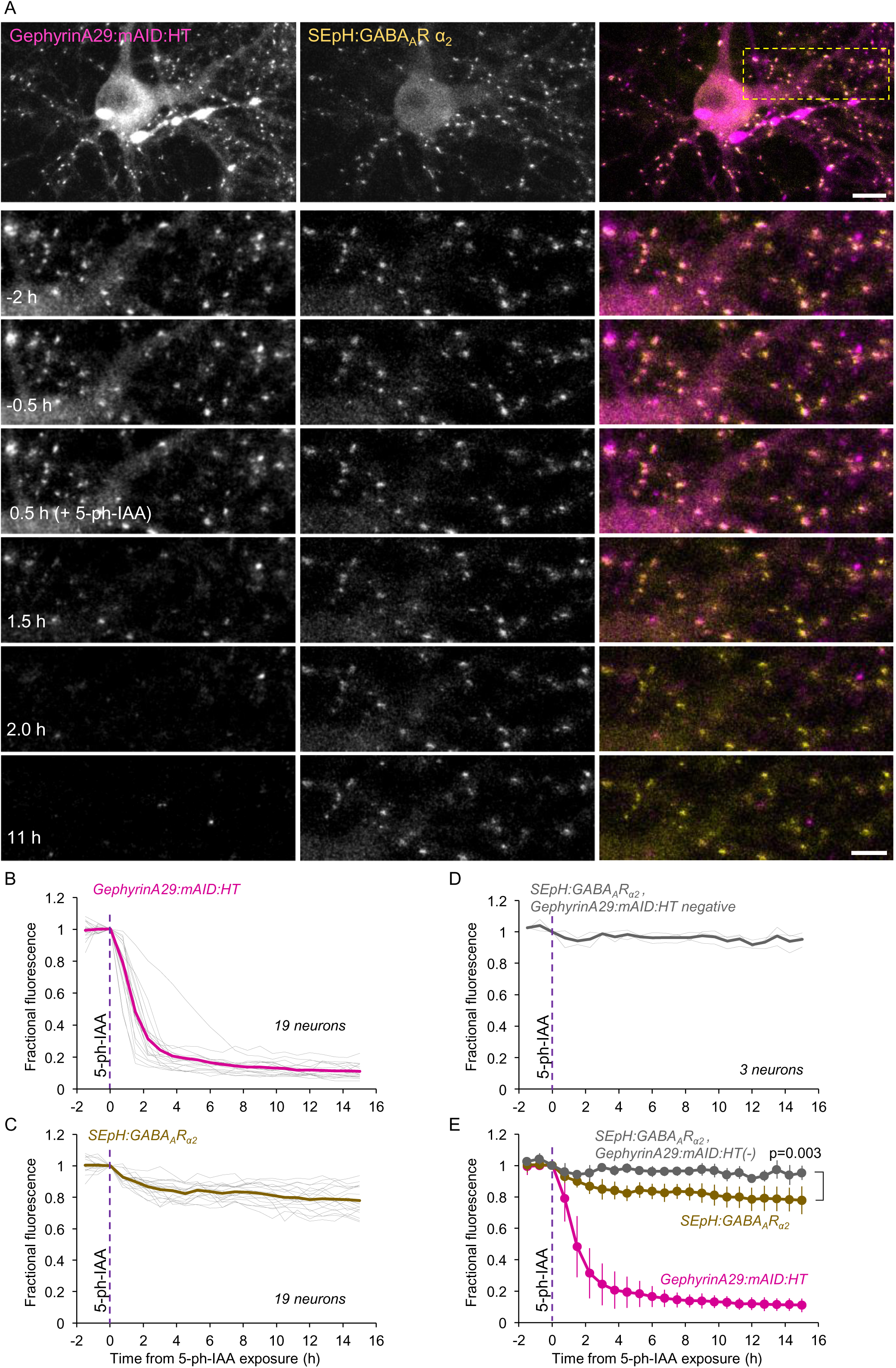
Acute elimination of GephyrinA29:mAID:HT is followed by loss of GABARs at the same synapses. A) Top panels: A rat cortical neuron in culture co-expressing GephyrinA29:mAID:HT (labeled with JF635-HT), SEpH:GABA_A_Rα_2_ and OsTIR1-P2A-mCherry (not shown). Bottom panels: Region in yellow rectangle at greater detail, before, and after addition of 5-Ph-IAA. Note that the near complete loss of JF635-HT fluorescence (presumably reflecting GephyrinA29:mAID:HT degradation) is associated with a modest reduction in SEpH:GABA_A_Rα_2_ fluorescence. Scale bars: 10 µm (top panels) 5µm (bottom panels). B) JF635-HT fluorescence measured at 50 synapses of each neuron tracked throughout the experiments. Each thin gray line is the average JF635-HT fluorescence measured for the synapses of one neuron (18 neurons from 3 experiments). Thick magenta line is the population average**. C)** changes in SEpH:GABA_A_Rα_2_ fluorescence at the same synapses and neurons of B. Thick brown line is the population average. **D)** SEpH:GABA_A_Rα_2_ fluorescence measured at 150 synapses of neurons positive for SEpH:GABA_A_Rα_2_ and OsTIR1-P2A-mCherry but negative for GephyrinA29:mAID:HT. Each thin gray line is the average fluorescence measured for the synapses of one neuron (3 neurons from 2 experiments). Thick gray line is the population average. **E)** Pooled data. Error bars are standard deviations. Test for difference between GephyrinA29:mAID:HT positive and negative cells – unpaired t-test, without assuming equal variances; applied to data of last time point.

The relatively modest loss of GABARs upon elimination of exogenous gephyrin indicated that the contributions of exogenous gephyrin to total synaptic gephyrin pools might have been less substantial than those of exogenous PSD-95. Indeed, comparing synaptic gephyrin levels in GephyrinA29:mAID:HT expressing and naïve neurons using quantitative immunocytochemistry, revealed a ∼1.8 fold increase in total gephyrin levels (16 fields of view in each condition, 2 separate experiments; Supp. Fig. 6), indicating a limited contribution of exogenous gephyrin to GABAR confinement. On a synapse by synapse basis we observed a modest positive correlation between the loss of SEpH:GABA_A_Rα_2_ and the loss of GephyrinA29:mAID:HT (Supp. Fig. 5E), which was lower than what was observed for excitatory synapses (Supp. Fig. 5A), possibly due to lower overexpression and the use of a HT label as compared to a fused fluorescent protein.

Similar experiments were carried out using the N-terminally tagged Gephyrin variant mAID:mTurq2:Gephyrin. mAID:mTurq2:Gephyrin and SEpH:GABA_A_Rα_2_ also colocalized extremely well, with a very high correlation of fluorescence at individual synapses (r= 0.86; p = 4.3*10^-268^; 900 synapses from 18 neurons from 3 experiments). The loss of mAID:mTurq2:Gephyrin was associated with a ∼40% reduction of SEpH:GABA_A_Rα_2_ fluorescence (Supp. Fig. 7A-D) which closely followed the time course of mAID:mTurq2:Gephyrin elimination.

These experiments demonstrate that the abrupt elimination of PSD-95 and Gephyrin results in a significant loss of AMPA and GABA_A_ receptors at the same synapses, respectively.

### GKAP elimination but not PSD-95 elimination reduces postsynaptic scaffold size

The ability to acutely degrade PSD-95 - a major PSD protein - created an opportunity to examine how its abrupt elimination affects other PSD proteins that collectively make up the PSD. As a proof of principle, we examined how PSD-95 loss affects GKAP, which has been shown to bind the guanylate kinase domain of PSD-95 (e.g. Kim et al., 1997; Zhu et al., 2017). To that end, we triple expressed PSD-95:mTurq2:mAID, OsTIR1-P2A-mCherry and a fusion protein of GKAP and mCitrine (mCit:GKAP) in cultured rat cortical neurons. Under baseline conditions, the correlation of mCit:GKAP and PSD-95:mTurq2:mAID fluorescence on a synapse-to-synapse basis was very high (r= 0.82; p = 7.6*10^-106^ 422 synapses from 17 cells in 3 experiments). As shown in Fig. 7, exposure to 5-Ph-IAA and the consequential loss of PSD-95:mTurq2:mAID were not associated with a concomitant loss of mCit:GKAP (see also Yao et al., 2003). Instead, and somewhat unexpectedly, its loss was accompanied by a ∼30% *increase* in mCit:GKAP content at individual synapses (12 neurons from 2 experiments). Comparisons with neurons in the same dishes expressing mCit:GKAP but not PSD-95:mTurq2:mAID revealed that the increase in mCit:GKAP fluorescence was highly significant (p = 1.0*10^-4^; 12 and 6 neurons, respectively). Similar observations were made in a separate set of experiments, using synapses identified automatically at each time point, rather than individually tracked synapses (Supp. Fig. 8; 13 neurons and 9 control neurons, from 3 experiments, >8,000 synapses per time point). These data indicate that in these experiments, PSD-95 was not the determining factor in setting the ‘size’ of these PSDs, and that its loss was followed by the ‘infiltration’ of, or substitution by other scaffold proteins, in particular GKAP. Indeed, when we co-expressed a fusion protein of PSD-95 and mCitrine (PSD-95:mCit) instead of mCit:GKAP, we observed a nearly identical phenomenon, namely the replacement and coalescence of cytosolic PSD-95:mCit at sites vacated of PSD-95:mTurq2:mAID (Supp. Fig. 9). Interestingly, the correlation at individual synapses of PSD-95:mTurq2:mAID and PSD-95:mCit fluorescence – identical proteins that differ only in the GFP variant they are fused to – was nearly identical (r= 0.82; p = 4.6*10^-114^; 450 synapses from 22 neurons in 3 experiments) indicating that this value is near the ceiling of such measurements in our experiments.

**Figure 7.**
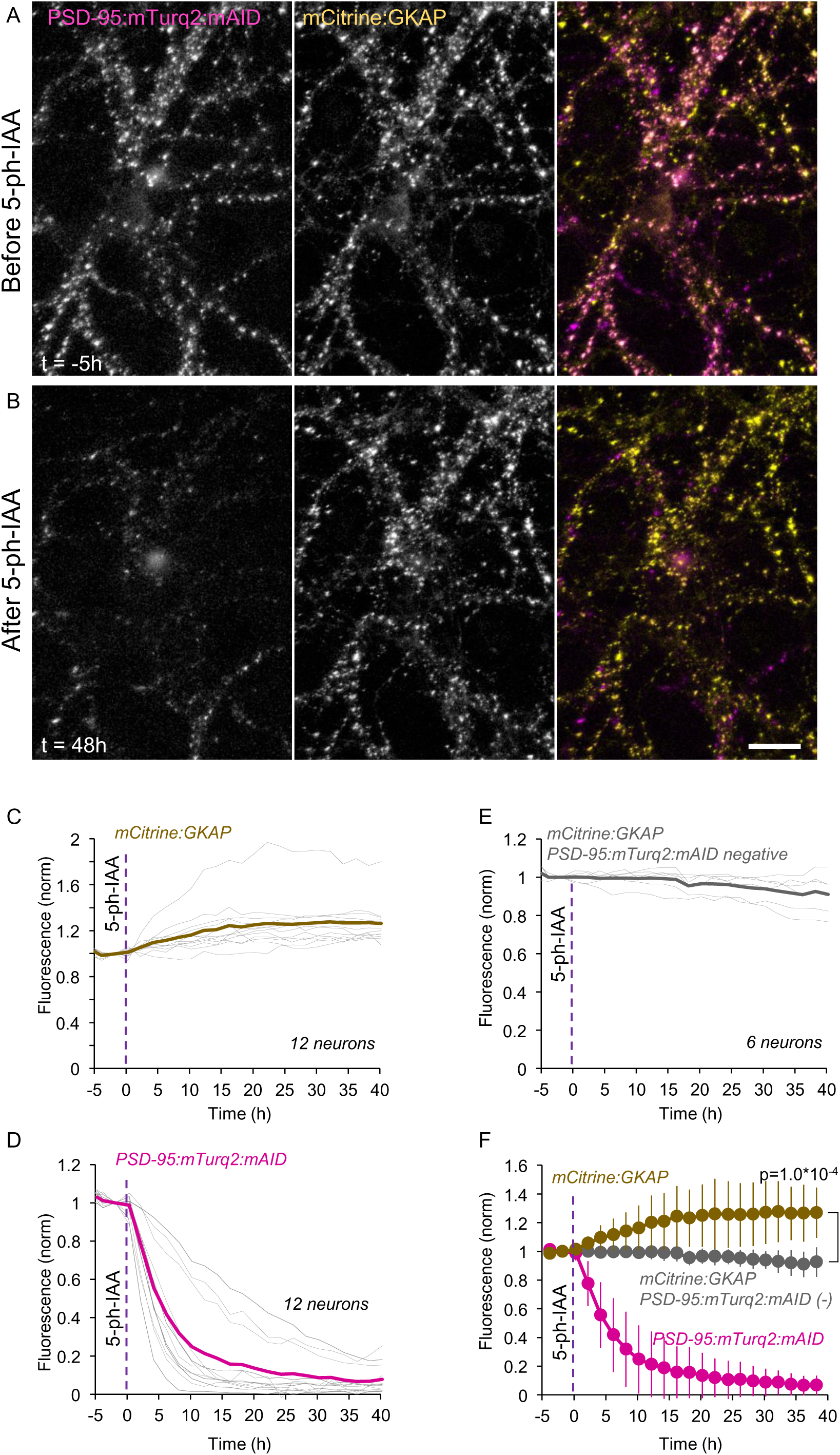
Acute elimination of PSD-95:mTurq2:mAID is followed by ‘substitution’ with GKAP. **A)** A rat cortical neuron in culture co-expressing PSD-95:mTurq2:mAID (left) and mCit:GKAP (middle) as well as OsTIR1-P2A-mCherry (not shown). **B)** 5-Ph-IAA induced nearly complete loss of PSD-95:mTurq2:mAID, and increased synaptic levels of mCit:GKAP at the same synapses. Scale bar: 20 µm. **C)** Changes in mCit:GKAP fluorescence measured at 13-40 synapses of each neuron tracked throughout the experiments (303 in total). Each thin gray line is the average normalized fluorescence measured for the synapses of one neuron (12 neurons from 2 experiments). Thick brown line is the population average. **D)** PSD-95:mTurq2:mAID fluorescence measured at the same synapses and neurons of C. Thick magenta line is the population average. **E)** mCit:GKAP fluorescence measured at 18-33 synapses of neurons positive for mCit:GKAP and OsTIR1-P2A-mCherry but negative for PSD-95:mTurq2:mAID (147 in total). Each thin gray line is the average fluorescence measured for the synapses of one neuron (6 neurons from 2 experiments). Thick gray line is the population average. **F)** Pooled data. Error bars are standard deviations. Test for difference between PSD-95:mTurq2:mAID positive and negative cells – unpaired t-test, without assuming equal variances; applied to data of last time point. See also Supp. Fig. 8.

Although PSD-95 is commonly viewed as a central organizer of PSD size and properties, other studies characterized GKAP as an equally important postsynaptic protein in this regard (Shin et al., 2012; Zhu et al., 2017). In fact, biochemical reconstitution studies using purified PSD proteins pointed to the singular importance of GKAP in driving PSD formation through Liquid-Liquid Phase Separation (LLPS; Zeng et al., 2018). We thus performed the reverse experiment, that is, rapidly eliminated GKAP and followed the consequences to PSD-95 contents at the same synapses. To that end we triple expressed mAID:mTurq2:GKAP, PSD-95:mCit and OsTIR1-P2A-mCherry. Here too, the correlation of mAID:mTurq2:GKAP and PSD-95:mCit fluorescence at individual synapses was maximal (r= 0.85; p = 5*10^-132^ 478 synapses from 21 cells in 3 experiments). Unlike what was observed for PSD-95, acute degradation of mAID:mTurq2:GKAP was associated with a marked reduction (∼30%) of PSD-95:mCit contents at the same synapses, which closely followed the time course of mAID:mTurq2:GKAP loss (Fig. 8A-E,H; 40 neurons from 5 experiments, p=9.5*10^-18^ when compared to 18 neurons that expressed PSD-95:mCit but not mAID:mTurq2:GKAP). Strikingly, the loss of synaptically associated PSD-95:mCit was associated with a parallel increase in cytosolic PSD-95:mCit levels (by ∼35% when measured at the cell body; Fig. 8 A,B, F-H; 31 neurons from 5 experiments, p=1*10^-8^ when compared to 14 neurons that expressed PSD-95:mCit but not mAID:mTurq2:GKAP). Thus, and unlike the effects of PSD-95 elimination, acute GKAP elimination caused the dissociation of PSD-95 from postsynaptic densities and apparent PSD ‘shrinkage’.

**Figure 8.**
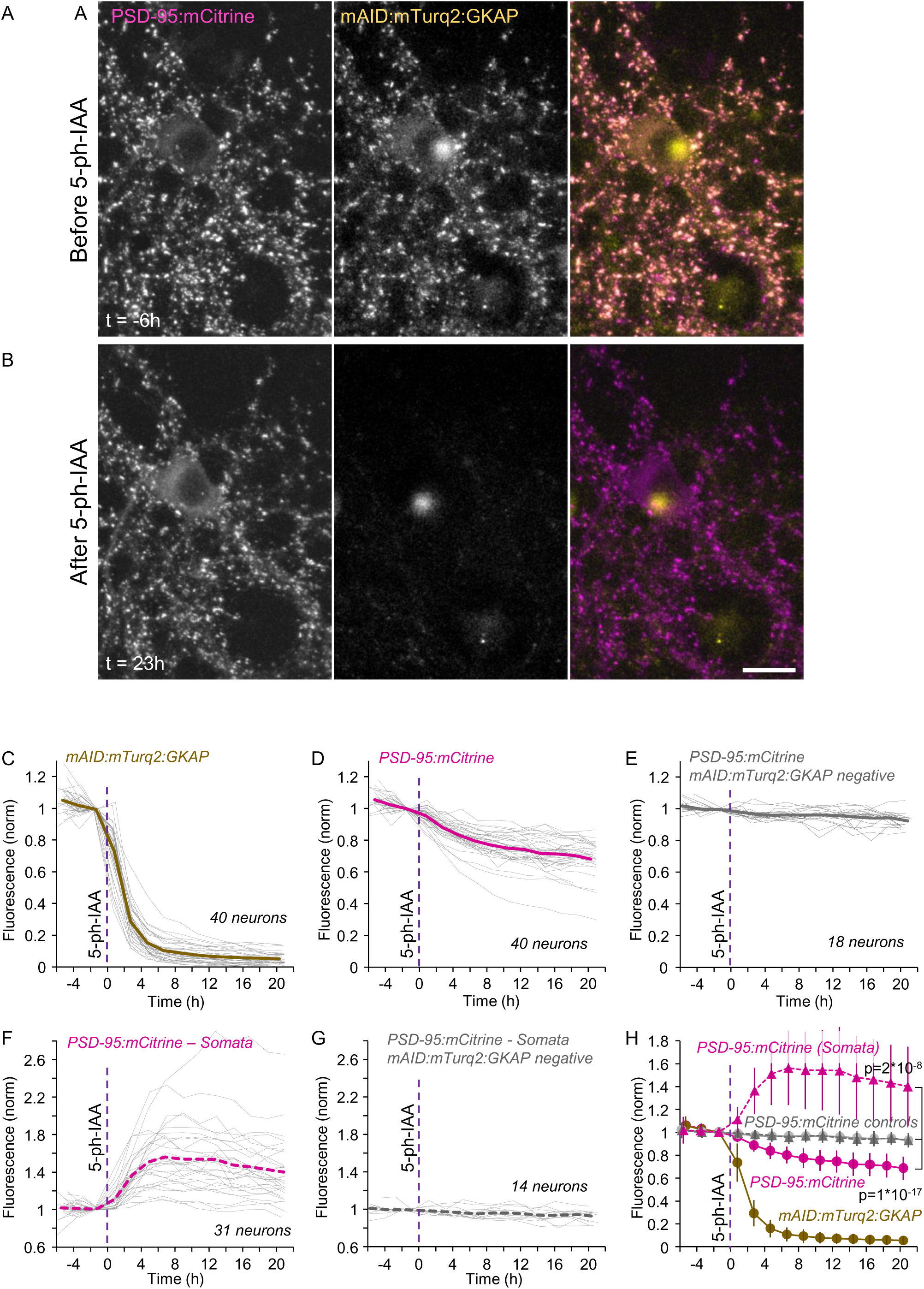
Acute elimination of mAID:mTurq2:GKAP is followed by concomitant loss of synaptic PSD-95 and elevated levels of cytosolic PSD-95. **A)** A rat cortical neuron in culture co-expressing PSD-95:mCit (left) mAID:mTurq2:GKAP (middle) as well as OsTIR1-P2A-mCherry (not shown). **B)** 5-Ph-IAA induced nearly complete loss of mAID:mTurq2:GKAP, loss of PSD-95:mCit from the same synapses and elevated levels of cytosolic PSD-95:mCit. Scale bar: 20 µm. **C)** Changes in mAID:mTurq2:GKAP fluorescence measured at 24-78 synapses of each neuron tracked throughout the experiments (1,723 synapses in total). Each thin gray line is the average normalized fluorescence measured for the synapses of one neuron (40 neurons from 5 experiments). Thick brown line is the population average. **D)** PSD-95:mCit fluorescence measured at the same synapses and neurons of C. Thick magenta line is the population average. **E)** PSD-95:mCit fluorescence measured at 21-64 synapses of neurons positive for PSD-95:mCit and OsTIR1-P2A-mCherry but negative for mAID:mTurq2:GKAP (752 synapses in total). Each thin gray line is the average fluorescence measured for the synapses of one neuron (18 neurons from 5 experiments). Thick gray line is the population average. **F)** Changes in cytosolic levels of PSD-95:mCit measured at the cell soma (31 neurons from 5 experiments). Dashed thick magenta line is the population average. **G)** Changes in cytosolic PSD-95:mCit measured in the cell bodies of neurons positive for PSD-95:mCit and OsTIR1-P2A-mCherry but negative for mAID:mTurq2:GKAP (14 neurons from 5 experiments). Dashed thick gray line is the population average. **H)** Pooled data. Dashed lines with filled triangles represent cytosolic PSD-95:mCit levels. Error bars are standard deviations. Tests for difference between PSD-95:mTurq2:mAID positive and negative cells – unpaired t-tests, without assuming equal variances; applied to data of last time points.

Comparisons on a synapse to synapse basis revealed a positive correlation between the loss of mAID:mTurq2:GKAP and PSD-95:mCit fluorescence after 15 h of exposure to 5-Ph-IAA (r = 0.41, 1,723 synapses from 39 neurons from 5 experiments; correlations determined for each neuron separately: 0.43 ± 0.17; mean ± standard deviation, respectively). Here too, PSD-95:mCit fluorescence loss was quite variable, with some synapses exhibiting no loss and even some gain (Supp. Fig. 5F), possibly reflecting some PSD remodeling that occurred during this time frame (e.g. Dvorkin and Ziv, 2016).

These experiments demonstrate how acute, mAID mediated degradation can be used to examine the roles of individual synaptic proteins in PSD organization.

## Discussion

We set out to examine the utility of the AID2 system as a tool for efficient, rapid, and reversible elimination of key synaptic proteins and for studying corresponding real-time consequences of acute protein elimination at individual synapses. We first established that exogenous fusion proteins of PSD-95, GKAP, and gephyrin N-terminally, C-terminally, or intramolecularly tagged with a minimal auxin induced degron (mAID) in conjunction with a fluorescent protein (mTurq2) or a HaloTag protein, localize correctly to synapses, and are rapidly degraded upon exposure to sub-micromolar concentrations of a small, membrane permeable inducer. Degradation was rapid and nearly complete, yet reversible upon removal of the inducer. We further established that the AID2 technology also allows for the rapid depletion of cytosolic and synaptic proteins *in vivo*, with kinetics comparable to those observed in cell culture. The rapid elimination of the corresponding proteins is associated with detectable cell biological effects, some expected, like receptor loss, and others less so, like PSD composition. Altogether, these findings show that AID2 technology is a uniquely powerful tool for studying synapse biology. Furthermore, the rapid onset, speed, and inherent reversibility of protein degradation offer many advantages in comparison to current approaches and thus provide means for performing stringent subtractive biochemistry experiments in living tissues, cells, and subcellular compartments, such as synapses.

### Advantages and limitations

The main advantage of AID2 technology in comparison to common knock-out and knock-down approaches is found in its capacity to eliminate a POI rapidly, bypassing the slow degradation pathways that knock-out and knock-down approaches depend on. The dramatic acceleration of POI elimination times from days to hours creates time windows within which direct consequences of POI elimination can be studied with minimal ‘contamination’ from slow adjustment/compensation processes that often complicate interpretations (e.g. Elias et al, 2006; Levy et al., 2015). The shortening of degradation times also has practical advantages in the sense that it greatly reduces the technical challenges associated with following POI elimination and consequential phenomena longitudinally in real-time, thus facilitating the application of this technology to numerous research questions and experimental systems.

A second advantage lies in AID2 simplicity: While other systems have been developed for rapidly degrading POIs, including ProTACs, molecular glue degraders (Tsai et al., 2024), GFE3 (Gross et al., 2016) and peptide-directed lysosomal degradation (Fan et al., 2014), these typically involve specific molecules tuned to target specific POIs. In contrast, AID2 involves a generic degron that can be used to target a diverse set of proteins, with our data indicating that degron location along the polypeptide chain is quite flexible.

A third advantage is its inherent reversibility. In this respect, new genetic approaches (e.g. Shi et al., 2022) also offer reversable gene manipulation. Yet, the simplicity of the AID2 system induction (and termination) mechanism - a small molecule that is easily introduced and removed or cleared, both in culture and *in vivo*, would seem advantageous compared to the complex molecular machineries of reversable genetic approaches.

A final, although not exclusive, advantage is the systems three component composition, with each component potentially controlling a separate experimental dimension: the mAID fusion protein (what), OsTIR1 (where) and 5-Ph-IAA (when).

The main disadvantage of the AID2 system its dependence on an exogenous tag fused to the POI, a modification with unexpected, potentially detrimental consequences. This is particularly true when generating knock-in animals in which degrons are inserted into genomic loci (see below). In simpler settings, the fusion protein is expressed on a background of endogenous variants of the same protein, an additional confound that must be considered. Yet, overexpression of exogenous POIs or protein fragments is a very common experimental manipulation in many settings. Here, the ability to reversibly control overexpression levels can be highly desirable as a means to examine effects in the same cells, tissues or animals over time. For example, such overexpression, once effected, can be terminated rapidly or titrated downwards at will, to examine the consequences of removing or pausing the experimental manipulation. Conversely, expression can be constitutively suppressed by chronic exposure to 5-Ph-IAA until overexpression is desired. Then, expression can be suppressed again by reintroducing the small ligand. In fact, cyclic overexpression also becomes possible.

Our study demonstrates that AID2 technology can be readily used *in-vivo* to transiently deplete cytosolic and synaptic target proteins, and that POI elimination and recovery can be followed in the same neurons over time. By way of extension, the optimal manner to use this approach and avoid the aforementioned overexpression issues would be to create knock-in animals in which the degron is added to the genomic locus of a POI, such that all of the POI contains the small degron. By crossing such animals with transgenic animals which express OsTIR (F74G) from cell-specific promotors (in analogy to the many CRE animals made to date), rapid and reversable elimination of a POI in specific cells, brain regions or organs could be achieved at will. Indeed, this possibility was very recently demonstrated in a proof-of-principle study (Makino-Itou et al, 2024) in which transgenic mice were made that express OsTIR1 specifically in germ cells, leading to selective degradation of mAID fusion proteins in these cells only. As an alternative, OsTIR (F74G) could be expressed locally (constitutively or cell-specifically) using viral expression vectors with appropriate promoters.

### *In situ* consequences of acute synaptic POI elimination

The acute and rapid elimination of mAID-tagged variants of the major synaptic scaffold proteins PSD-95 and Gephyrin had clearly detectable and defined cell biological consequences, even though the endogenous variants of the two proteins were not explicitly manipulated. Given their known roles, the consequential loss of AMPA and GABA receptors, respectively (Figs. 5,6, Supp. Fig. 7), was expected. In fact, the findings are in line with prior studies using photoinactivation of PSD-95 (Yudowski et al., 2013) and elimination of gephyrin using a fusion protein of an E3 ligase and an antibody-like protein (FingR) targeted at gephyrin (Gross et al., 2016). However, in comparison to the nearly complete loss of the tagged proteins, receptor loss was relatively modest. This might indicate that receptor sequestration by endogenous scaffold proteins is already quite close to saturation with the endogenous levels of scaffolding proteins (but see, e.g., El-Husseini et al., 2000; Futai et al., 2007; Kim et al., 2007, Béïque and Andrade, 2003; Ehrlich and Malinow, 2004; Elias et al., 2006; Ehrlich et al., 2007; Schlüter et al., 2006). Alternatively, the degeneracy of PSD proteins, in particular at excitatory synapses (e.g. Elias et al., 2006; Levy et al., 2015), might limit the impact of removing a single PSD scaffold protein species on receptor sequestration at postsynaptic membranes. Indeed, genetic elimination of PSD-95 perturbs, but does not abolish AMPAR mediated postsynaptic currents (e.g. Migaud et al., 1998; Béïque et al., 2006). It is also possible that expression levels of receptor subunits or auxiliary proteins involved in receptor retention might not scale to match those of the exogenous scaffold proteins. Finally, it cannot be ruled out that the fluorescent tags and/or the exogenous degron affect scaffold protein function, which might have been the case for mAID:mTurq2:Gephyrin. We note, however, that knock-in mice expressing PSD-95-EGFP (Zhu et al., 2018) and PSD-95-HaloTag (Bulovaite et al., 2022) as well as Gephyrin - RFP (Machado et al., 2011) fusion proteins, very similar to those used here, are viable without overt phenotypical defects.

The observation that the acute loss of PSD-95 was associated with parallel increases in GKAP and degron-free PSD-95:mCit levels at the same synapses (Fig. 7, Supp Fig. 8) was less expected. The most likely explanation is that scaffold size is determined collectively by a large number of synaptic protein species and that the loss of one of these does not necessarily reduce scaffold size, only its structure and composition, creating ‘openings’ for the binding of other proteins (see for example Chen et al., 2011). Indeed, the complete knock out of PSD-95 does not impact dendritic spine volume in the mouse hippocampus (Béïque et al., 2006). Our findings, and in particular the opposing effects of PSD-95 and GKAP elimination, are especially intriguing when considered within a framework positing that PSDs act as molecular condensates formed through LLPS, (e.g. Zeng et al., 2018, 2019; Bai et al., 2021; Vistrup-Parry et al., 2021; Zhu et al., 2024; reviewed in Chen et al., 2020; Hayashi et al., 2021; Bai and Zhang 2022). Evidence for this framework comes primarily from biochemical reconstitution experiments, in which the formation of visible condensates was quantified following the mixing of purified PSD proteins. Particularly noteworthy in the context of the current findings is the study of Zeng et al., 2018, in which condensate formation in mixtures of purified PSD-95, GKAP, Shank and homer was found to be uniquely dependent on the presence of GKAP. The similar observations made here (Figs. 7, 8) highlight the potential of the AID2 system to carry out analogous experiments in living cells, to identify proteins that act as ‘drivers’ (i.e. induce condensate formation) or join as relatively passively as ‘clients’ (Zeng et al., 2018).

In sum, our study illustrates the utility of the AID2 system to decipher molecular mechanisms that sculpt synapses, their contents and functions, and the roles played by individual proteins in these processes. As noted above, however, this system also creates exciting opportunities for studying “chemistry *in cellulo*” (Salvatico et al., 2015), and as such, is applicable to both focused and general questions concerning specificity and degeneracy in networks of interacting proteins.

## Methods

### Animals and cell culture preparations

Primary cultures of cortical neurons were prepared from newborn Wistar rats (either sex), following protocols approved by the Technion Israel Institute of Technology’s Committee for the Supervision of Animal Experiments (approval IL-105-08-20). In brief, cortices from 0 to 1-day-old rats were dissected and dissociated using trypsin, followed by gentle trituration with a siliconized Pasteur pipette. Approximately 5 × 10^5^ of dissociated cells were then plated on polyethylenimine-coated 29 mm glass-bottom dishes (MatTek) to promote adherence. The neurons were initially grown in a medium consisting of Eagle’s minimal essential medium (Sigma-Aldrich), 25 µg/mL insulin (Sigma-Aldrich), 20 mM glucose (Sigma-Aldrich), 2 mM L-glutamine (Sigma-Aldrich), and 10% NuSerum (Becton Dickinson Labware). Cultures were maintained in a humidified incubator at 37°C with 95% air and 5% CO_2_. From day 7, half of the culture media was replaced three times a week with feeding medium, similar to the media described above, except for the absence of NuSerum, a reduced concentration of L-glutamine (0.5 mM), and the addition of 2% B-27 supplement (Gibco).

For the *in-vivo* experiments (Fig. 3 and 4), wildtype C57BL/6J mice were used. At the time of starting the experiments, all animals were at ages of six to eight weeks. For the experiments of Fig. 3 a total of 3 mice were used (each iteratively underwent multiple treatment conditions). For the experiment of Fig. 4, 7 mice were used that were split into two groups receiving different treatment (n = 4, 5-Ph-IAA, n = 3 sham treatment). All animal experiments were performed in accordance with the German laboratory animal law guidelines for animal research and had been approved by the Landesuntersuchungsamt Rheinland Pfalz (Approval # G 17-1-051 and G 22-1-091).

### DNA constructs

For experiments carried out in cell culture, fusion proteins were introduced using third-generation lentiviral expression vectors based on a modified FUGW (FUGWm) backbone (Lois et al., 2002) in which an XhoI restriction site was moved to a downstream position. Full, annotated sequences of all lentiviral plasmids used here (.seq files) are provided as supplementary materials. All gene synthesis and cloning were done by Genscript (Piscataway NJ, USA).

Lentiviral vectors for expressing OsTIR1(F74G) in neurons in culture were created as follows: The pAAV-hSyn-OsTIR1(F74G) plasmid described in Yesbolatova et al., 2020 was obtained from Addgene (Addgene #140730). Then, the Synapsin promotor and the coding region were cut out of pAAV-hSyn-OsTIR1-F74G with MluI and BclI. PacI (5’) and XhoI (3’) sites were added to the excised segment which was then inserted into FUGWm at its PacI and XhoI sites, resulting in OsTIR1-P2A-EGFP:mAID. OsTIR1-P2A-mCherry was created by full length synthesis of mCherry flanked by BsmBI and BstBI and replacing the mAID:EGFP segment in OsTIR1-P2A-EGFP:mAID with this insert.

The vector encoding for SEpH:GluA2 was described previously (Zeidan et al., 2012). All other plasmids used here (PSD-95:mCit, mCit:GKAP, PSD95:mTurq2:mAID, mAID:mTurq2:GKAP, SEpH:GABA_A_Rα2, mAID:mTurq2:Gephyrin, PSD-95:HT:mAID, GephyrinA29:mAID:HT) were made by large scale gene synthesis of the POI followed by insertion into the modified FUGWm backbone described above using appropriate restriction sites. All inserts as well as at least 200 flanking base pairs were sequenced and checked for correctness.

### Lentivirus production and transduction

Lentiviral particles were generated by transfecting HEK293T cells with a plasmid mixture containing essential HIV packaging genes and a heterologous viral envelope gene (MISSION® Lentiviral Packaging Mix, Sigma). Transfection was carried out using Lipofectamine 2000 (Invitrogen), with HEK293T cells raised on 10 cm plates at approximately 90% confluence. The supernatant was collected 48–72 h post-transfection, filtered through 0.45μm filters, aliquoted, and stored at −80°C. Neurons were infected with either one or more of the constructs mentioned earlier. For double or triple infections, the viral particles were mixed before they were added to the plates. The infection was done for most experiments on 9-10 post-plating, except for the experiments of the triple expression of mAID-mTurq2-GKAP1 with PSD-95:mCit and OsTIR1-P2A-mCherry in which the neurons were infected on day 3 post-plating.

### AAV production

For AAV production 6 × 10^7^ HEK 293 cells were seeded in a 16-layer Celldisc (Greiner; Cat. no. 678916) with 1L complete growth media (DMEM, Gibco; Cat. No. 52100–047), supplemented with 10% heat-inactivated FBS (Sigma; Cat. No. F7524), 2 mM L-glutamine (Sigma; cat. no. G7513) and 1% Penicillin Streptomycin (Sigma-Aldrich Cat. No. P0781-100ML) and cultured for 48 h in CO_2_ incubator (37 °C temperature, 95% relative humidity and 5% CO_2_). For chemical transfection plasmid pADDeltaF6 (Addgene Cat. No. #112867), pAAV8 (Addgene Cat. No. # 112864) and the respective expression vector plasmid were mixed at equimolar ratio to a total of 2.069 mg DNA. 69 ml of 300 mM CaCl_2_ was added to the plasmid DNA. The entire CaCl_2_/DNA mixture was slowly added to 69 ml 2xHBS solution (Aesar; Cat. No. #J62623). After 5 min. incubation the mixture was added to 500 ml DEMEM supplemented with 5% FCS (no antibiotics). Culture media was then carefully decanted from the Celldisc and replaced with the transfection media. After 6 h incubation (37 °C, 95% relative humidity and 5% CO_2_) transfection media was carefully decanted and replaced with 1L of fresh complete growth media. Transfected cells were incubated for 72 h (37 °C, 95% relative humidity and 5% CO_2_).

To harvest the cells growth media was carefully decanted and collected. 500 ml of kept growth media was supplemented with 7 ml 0.5 M EDTA (Invitrogen; Cat. No. #15575-020) and 400 ml out of it was put back into the Celldisc. After 5 min incubation at room temperature cells detached from the surface. Cell suspension was transferred to a 500 ml centrifugation flask (Corning; Cat. No. 431123). The remaining 100 ml Growth media/EDTA mix was used to wash the Celldisc and added to the centrifugation flask.

After centrifugation at 800 × g for 15 min at 4 °C, supernatant was carefully discarded. The cell pellet was resuspended in 10 ml PBS, transferred to a 50 ml Falcon tube and centrifuged again for 15 min at 800 × g, at 4 °C. PBS was then discarded and the pellet resuspended in 24 ml lysis buffer (50 mM Tris, 1 M NaCl, 10 mM MgCL2) supplemented with 0,001% Pluronic F-68: (Invitrogen #24040032), 1300U Salt Active Nuclease (SAN; Sarstedt #83.1803) and 100 × HALT Protease Inhibitor Cocktail, (EDTA-free Thermo scientific #78439). Cell suspension was then subjected to three freeze / thaw cycles in liquid nitrogen and a 37 °C water bath, respectively. To assure that the suspension does not contain any remaining plasmid DNA, it was again supplemented with 1300U of SAN afterwards, and incubated at 37 °C for 1 h, while shaking at 150 rpm. Following centrifugation at 2500 × g for 15 min at r.t, cell debris was discarded, and the supernatant was transferred to a new 50 ml Falcon tube. 40% PEG-8000 solution (Polyethyleneglycol, Sigma #89510, in H2O, supplemented with 0.001% Pluronic) was added to a final concentration of 8%, mixed and incubated on ice at 4 °C for 16 h to 24 h. After centrifugation at 2500 × g for 30 min at 4 °C, supernatant was discarded, any residual PEG was carefully removed and 14.5 ml resuspension buffer (50 mM TRIS, 1 M NaCl, 0.001% Pluronic, pH8.0) was added to the pellet and the pellet was resuspended by vortexing and pipetting before it was incubated for at least 24 h at 4 °C, while shaking at 350–400 rpm. It was found crucial to resuspend the pellet completely. The suspension was then centrifuged at 2500 × g for 30 min at 4 °C and the supernatant was transferred to an ultra-centrifugation tube (Quickseal Tubes, Beckman Coulter #342414). AAV purification was performed by ultra-centrifugation over a discontinuous Iodixanol density gradient (OptiPrep Density Gradient Medium, Sigma #D-1556, 60% solution in H2O), as described previously with Iodixanol phases of 15%, 25%, 40% and 54%, respectively. After centrifugation, approximately 3.5 ml of the Iodixanol phases containing the filled AAV capsids were collected (2.5 ml of 40% and 1 ml of 54% phase). Special care was taken not to touch the 25% phase, since it contains empty capsids. For buffer exchange and concentration, AAV purification buffer (1 × PBS, 1 mM MgCl2, 2.5 mM KCL, 0.001% Pluronic, pH 7.4) was added to the virus containing fraction, to a total volume of 12 ml and transferred to a 15 ml AMICON ULTRA-15 column; (MWCO 100 kDa, Millipore #UFC910024). Centrifugation was performed according to the manufacturers protocol. After concentration of the virus solution to approximately 1.5 ml, fresh AAV purification buffer was added to a total volume of 12 ml, and centrifugation was repeated. At a volume of 0.5 ml to 1 ml, virus solution was resuspended thoroughly, transferred to a new tube and stored at -80 °C. The genomic titer was determined by qRT-PCR.

### Stereotaxic injection

All surgical equipment was sterilized with 70% ethanol before use. Animals were deeply anesthetized with isofluorane (Abbott Animal Health, IL, USA; IsoFlo) and positioned in a stereotaxic frame (Kopf Instruments, Tujunga, CA, USA; Stereotaxic System Kopf 1900). The eyes were protected from dehydration and intensive light exposure using Vaseline and a piece of aluminum foil. The anesthesia was maintained by delivery of a 1.5 to 2.4% isoflurane/air mixture with a vaporizer (High Precision Instruments, MT; Univentor 400 Anaesthesia Unit) at a flow rate of around 200 ml/min to the snout. Lidocaine was applied as local anesthetic subcutaneously before exposure of the skull. The scalp was washed with a 70% ethanol solution and a cut along the midline revealed the skull. A small hole was drilled into the skull above the auditory cortex using a motorized dental drill, leaving the dura mater intact. Injections were performed perpendicular to the surface of the skull. Virus solutions were specific for each experiment and consisted of a mixture of different AAV viruses in PBS: For the *in-vivo* experiments of Fig. 3, a 1:1 mix of AAV8-hSyn-OsTIR1(F74G)-P2A-mAID:EGFP:NES (plasmid from Yesbolatova et al., 2020, Addgene #140730, titer: 1.4 ∗ 10^15^ VG/mL) and AAV8-phSyn-H2B::mCherry (titer: 1.8 ∗ 10^14^ VG/mL) was injected. For the *in-vivo* experiments of Fig. 4, a 1:1:1 triple-mix of AAV8-hSyn-PSD-95:HT:mAID (approx. titer: 10^13^VG/mL), AAV8-hSyn-OsTIR1-P2A-NLS:tagBFP2 (approx. titer: 10^13^VG/mL) and AAV8-hSyn-PSD-95.FingR:EGFP-CCR5TC (titer: 1,0 ∗ 10^15^VG/mL) was injected. The virus mixture was loaded into a thin glass pipette and 170 nl were injected at a flow rate of 20 nl/min (World Precision Instruments, Sarasota, FL, USA; Nanoliter 2000 Injector) in five locations of the right auditory cortex, resulting in a total injection volume of 850 nl. Stereotactic coordinates were: 4.4, -2,5/-2.75/-3/-3.25/-3.5, 2.5 (in mm, lateral, caudal, and ventral in reference to Bregma). Glass pipettes (World Precision Instruments, Sarasota, FL, USA; Glass Capillaries for Nanoliter 2000; Order# 4878) had been pulled with a long taper and the tip was cut to a diameter of 20 to 40 μm. After the injection, the pipette was left in place for three minutes, before being slowly withdrawn and moved to the next coordinate. After completion of the injection protocol, the skin wound was sealed using tissue adhesive (3M Animal Care Products, St. Paul, MN, USA; 3M Vetbond Tissue Adhesive), and anesthesia was terminated. Mice were monitored daily and intraperitoneal injections of carprofen (0.2 ml of 0.5 mg/ml stock) were applied on the first days after surgery.

### Cranial window implantation

Four to eight weeks after stereotactic injections, animals were anesthetized using isoflurane (Abbott Animal Health, IL, USA; IsoFlo). All surgical equipment and glass cover slip were sterilized with 70% ethanol before use. Anesthesia was initialized in a glass desiccator filled with an isoflurane/air mixture. Anesthetized animals were mounted on a stereotaxic frame (Kopf Instruments, Tujunga, CA, USA; Stereotaxic System Kopf 1900) and the head was positioned using ear, teeth, and a custom-made V-shaped head holder. Anesthesia was maintained by delivery of a 1.5 to 2.4% isoflurane/air mixture with a vaporizer (High Precision Instruments, MT; Univentor 400 Anaesthesia Unit) at a flow rate of around 200 ml/min to the snout. 0.02 ml dexamethasone (4 mg/ml) was administered intramuscularly to the quadriceps, as well as 0.02 ml carprofen (0.5 mg/ml) intraperitoneally. The eyes were protected from dehydration and intensive light exposure using Vaseline and a piece of aluminum foil. Lidocaine was applied subcutaneously before exposure of the skull. The scalp was washed with a 70% ethanol in water solution and a flap of skin covering temporal, both parietal regions and part of the occipital bone was removed. All following surgery steps were conducted unilaterally on the right side of the animal: The temporal muscle was partly removed with a surgical scalpel and forceps to expose the right temporal bone. Using a fine motorized drill, the bones of the skull were smoothened, and part of the zygomatic process was removed. The surface was cleaned using cortex buffer and covered with a thin layer of one component-instant glue (Carl Roth, Germany; Roti coll). A thin layer of dental cement (Lang Dental, IL, USA; Ortho-Jet) was applied onto the skull, sparing the area of the temporal bone above the auditory cortex. An elliptic groove of about 3 mm diameter was carefully drilled into the skull above the auditory cortex, and the bone was carefully lifted using scalpel and forceps. The exposed area was carefully cleaned and kept moist using sterile sponge (Pfizer, NY, USA; Gelfoam) and cortex buffer. The craniotomy was covered with a small circular cover glass (Electron Microscopy Sciences, PA, USA; five mm diameter, catalogue #72195-05. The cover glass was finally set in place with one component-instant glue and dental cement. In order to position the animal under the microscope with the objective facing the window plane perpendicularly, a custom-made titanium head post was mounted on the implant above the window and embedded with dental cement. After dental cement had dried, animals were placed back in a pre-warmed cage. After the surgical procedure, animals recovered for at least one week before continuing the experiments.

### Confocal imaging (cell culture experiments)

Glass-bottom dishes containing cortical neurons expressing one or more of the previously mentioned fusion proteins were imaged on a custom-built confocal laser scanning (inverted) microscope based on the Zeiss Axio Observer Z1, equipped with a 40×, 1.3 N.A. Plan-Fluar objective. The system, controlled by custom software, was designed to allow automated, multisite time-lapse microscopy. Dishes were fitted with a custom-designed cap featuring inlet and outlet ports for perfusion and air exchange. Neurons were continuously perfused with feeding media (described above) starting 24 h before imaging commenced and continuing until the conclusion of the experiment. The perfusion feeding media was supplemented with 7-10% distilled water to compensate for evaporation and delivered to the dish at a flow rate of approximately 3 ml/day. This was achieved using custom perfusion systems based on ultraslow-flow peristaltic pumps (Instech Laboratories, Inc.) and silicone tubing, connected to the dish via the cap’s ports. A sterile gas mixture of 95% air / 5% CO_2_ was delivered into the dish at low rates, regulated by a high-precision flow meter (Gilmont Instruments), either directly to the dish through a dedicated inlet in the cap or using a closed chamber above the dish. The microscope’s objective and the base of the headstage were heated to 37°C using resistive elements with separate temperature sensors and controllers, maintaining the culture medium at 35–36°C.

Time-lapse imaging was typically performed by averaging five frames at 10-15 focal planes, spaced 0.8 µm apart. Images were captured at a resolution of 640×480 pixels, and 12 bits per pixel. Images were collected sequentially from multiple sites using a motorized stage that cycled automatically through locations at predetermined intervals. Focal drift was automatically corrected using the microscope’s autofocus system.

Fluorescent proteins and labels were imaged using the following excitation sources and emission filter combinations: mTurquoise2 – 457nm excitation (Argon laser, National Laser Company), FF01-475/20 or FF01 483/32 (Semrock), emission; EGFP and pHluorin – 488nm excitation (Argon laser), ET 525/50 (Chroma) or FF03 525/50 (Semrock), emission; mCitrine – 514nm excitation, (Argon laser), HQ545/50 (Chroma), emission; mCherry – 594nm excitation (DPSS laser; Cobolt), BLP01 594R (Semrock), emission; JF552-HT– 552nm excitation (DPSS laser; Coherent), ET 590/50 (Chroma), emission; JF635-HT and Alexa 647 / Cy5 decorated (secondary) antibodies – 632nm excitation (Helium Neon laser; JDS Uniphase/Thorlabs), RET 638 (Chroma) or LP02-633RU (Semrock), emission.

### Two-Photon Imaging (*in vivo* experiments)

The two-photon microscope (Prairie Technologies, WI, USA; Ultima IV) was comprised of a 20x-objective (Olympus, Tokyo, Japan; XLUMPlan Fl, NA = 0.95) and a pulsed laser (Coherent, CA, USA; Chameleon Ultra). All fluorophores (EGFP and mCherry) were co-excited at 920 nm wavelength, and separated by emission using a fluorescence filter cube (filter one: BP 480-550 nm; filter two: LP 590 nm; dichromatic mirror: DM 570 nm; Olympus, Tokyo, Japan; U-MSWG2). For the recordings, mice were lightly anaesthetized with isoflurane (flow from 0.9 to 1.6% isoflurane/air as described above) in order to avoid movement artifacts. Line scan imaging was performed using a field of view of 367x367μm (1024x1024 pixels) at a sampling period of 3.024 seconds per frame. For each mouse, and imaging session, one to three spots from the cortical surface were imaged, where for each spot a z-stack of 70 images was recorded with a step size of 2μm in between. For each spot, the z-level of the dura mater was identified and then, the image stack was recorded at a depth of 50-190 μm below the dura. For longitudinal tracking of the same cortical spots, the microscopic blood vessel patterns on the brain surface and the neuronal fluorescence patterns on superficial image planes were identified and matched to the reference recording from the first baseline time point. Animals were imaged on two baseline time points prior to any treatment (-1 and -0.5 h). Then, animals were randomly assigned to be injected with 10mg/kg 5-Ph-IAA or PBS. In the following, the same set of cortical spots were imaged repeatedly at intervals of 1, 3, 6, 9, 24, 48 and 72 h post injection. Between imaging periods, animals were placed back in their home cages. All mice repeatedly underwent both treatments (sham and 5-Ph-IAA) with intervals of approximately 1 week, where we ensured that before starting a novel recording round the mAID:EGFP fluorescence had fully recovered, matching the baseline conditions from the initial recording.

### 5-Ph-IAA Preparation and treatment

For cell culture experiments, 5-Ph-IAA (MedChemExpress, # HY-134653) was dissolved in DMSO and stored as 100mM aliquots at −80°C. For use, the 5-Ph-IAA was diluted in feeding medium to a final working concentration of 200nM.

For *in vivo* experiments, 5-Ph-IAA was dissolved in PBS. For the treatment, mice were injected with 120μl of the solution intraperitoneally, corresponding to 10mg/kg body weight. For the sham treatment, mice were intraperitoneally injected with 120μl PBS.

### HaloTag ligands preparation, labelling and staining

The Halotag ligands used in this study were Janelia Fluor® 635 HaloTag® Ligand (JF635-HT) and Janelia Fluor® 552 HaloTag® Ligand (JF552-HT), both were provided as generous gifts by Luke Lavis, Janelia Research Campus. (Grimm et al., 2017)

For the cell culture experiments, the ligands were dissolved in DMSO and kept as 100 µM aliquots stocks at −20 °C. The ligands were added to the dishes 1 h before the beginning of the experiment, at a final concentration of 100 nM. To assess the recovery of the tagged proteins following the wash, a second labelling with the ligands was performed.

For labeling brain slices, a stock solution of JF635-HT with DMSO, (D2650, Sigma Aldrich), Pluronic F-127, and sterile PBS (Mohar et al., 2022) was prepared. From this stock solution, a diluted working solution of 400nM JF635-HT in PBS was prepared just prior to addition to slices. This dye solution was added to the brain slices, followed by incubation at RT for 2 h. Finally, the dye was removed and the slices were washed two times for 15 minutes with PBS. Sections were mounted with Fluoromount-G.

### Preparation and imaging of histological samples

For the experiments of Fig. 4, mice were sacrificed via cervical dislocation 6 h after treatment with 5-Ph-IAA or PBS respectively. The brains were extracted, briefly cleaned in PBS and then transferred in 4% paraformaldehyde (PFA) in PBS. The brains stored in 4% PFA at 4°C for 24 h, before re-transferring them into PBS. Slices of 70 μm (coronal section) were obtained using a Leica VT1000S vibratome.

Images of the cortical regions were acquired with a 100x oil objective (Leica; HC PL APO CS2, NA = 1.4) using Stellaris 8 FALCON confocal microscope (Leica; funded by the Deutsche Forschungsgemeinschaft, project number 497669232). Field of views of 116.25x116.25μm (1664x1664 pixels) at layer2/3 of cortex are taken. Lasers 405, 488, and 640 were used to excite tagBFP, eGFP, and JF635 signals respectively. All channels were acquired in counting mode using line accumulations at speed of 400.

### Immunocytochemistry

Cultures of cortical neurons (naive or infected) were perfused with media for 24 hr as described above, imaged, and then fixed and stained as follows: Cells were washed with Tyrode’s physiological solution (119 mM NaCl, 2.5 mM KCl, 2 mM MgCl_2_, 25 mM HEPES; 30 mM Glucose and 2 mM CaCl_2_, pH 7.4) and exposed to a fixative solution (4% formaldehyde and 4% Sucrose in PBS) for 20 min at room temperature. This was followed by adding fixative solution supplemented with Triton X-100 (0.25%) for an additional 20 min, followed by washing with PBS. Fixed cells were then incubated with 10% bovine serum albumin (BSA) in water for 1 h at 37 °C. Following this blocking step, the cells were stained overnight at 4°C with FluoTag®-X2 anti-PSD95 Alexa 647 (NanoTag Biotechnologies # N3702-AF647-L; 1:200), or exposed to a mouse anti gephyrin primary antibody (Synaptic Systems anti gephyrin #147 111 or #147 011; 1:500 and 1:1000, respectively) overnight at 4 °C, followed by washing with PBS and labelling with a secondary antibody (Cy™5 AffiniPure conjugated polyclonal Donkey Anti-Mouse IgG; Jackson ImmunoResearch Laboratories, #715-225-150) (1:200) 1 hr at 37 °C. Cells expressing GephyrinA29:mAID:HT were labelled with the HT ligand JF552-HT prior to fixation.

### Image processing and analyses

All imaging data collected in cell culture were analysed using custom software (’OpenView’), which offers both automated and manual tracking of individual objects as well as measurements of fluorescent intensities over time. For fluorescence intensity measurements of synapses, regions of interest (ROIs; 9x9 pixels) were programmatically placed on fluorescent punctate objects at the initial time point and then semi-automatically tracked over time, focusing only on synapses that persisted throughout the experiment (synapses that split, morphed, merged, formed or disappeared were excluded from the analysis). Mean pixel intensities within these ROIs were calculated from maximal intensity projections of Z-stack sections. To assess cytosolic fluorescence levels, ROIs were manually positioned at initial time points over cell bodies, and mean pixel intensities were measured for each time point using Z-stack projections. Fluorescence values were corrected for background fluorescence by subtraction of values measured at cell-free regions, in some cases supplemented by spectral unmixing when fields of view contained substantial non-specific fluorescent objects.

In most cases, data collected from individual synapses were normalized by dividing fluorescence values collected from synapses (and somata) by values collected at the time point prior to the addition of 5-Ph-IAA.

Time constants for degradation curves were derived using a program written in Visual Basic for Applications within Microsoft Excel. The application attempts to fit the data to a sum of two exponentials, each with its own time constant, with a third parameter representing the relative fraction of each component. The application systematically explores a wide range of parameter and returns the values that result in a minimal sum of squared residuals. Time constants stated in the manuscript are for the large pools (78% to 98%). Time constants derived for minor pools were typically an order of magnitude longer, probably reflecting secondary processes such as photobleaching.

For the experiments of Fig. 3, the recorded image stacks from each cortical spot were fed into a custom standardized MATLAB processing pipeline (Muller thesis 2023; Eppler, 2022) that enabled us to track the same set of neurons from multiple defined image plains (syn. fields of view, FOV) of a stack and assess their cytosolic fluorescence signal over time (see Supp. Fig. 10A). (i) We first defined a selection of 13 FOV to analyze per stack. Specifically, we used the initial baseline recording to select the FOV on a z-level of 60μm below the dura mater as well as the following twelve deeper FOV in steps of 10μm. 1. (ii) For each recording time point, we identified the image planes that best matched these initial FOVs in respect to their nuclear H2B:mCherry signal. For this, we calculated key points on the images using a SIFT algorithm (scale-invariant feature transform) and cross-checked the key points with a brute force matching strategy (euclidean distance-based match-validation). (iii) We then automatically identified regions of interest from the H2B:mCherry images (ROIs corresponding to nuclei). For this, images were equalized with a Gaussian blur filter, binarized (Otsu threshold) and blob detection was applied, using a Laplacian-of-Gaussian algorithm. (iv) Each FOV was then aligned to the reference image from the initial time point, applying an xy-translation and tracking the ROIs with a local affine transformation. 2. (v) Lastly, we quantified the cytosolic fluorescence signal by defining a ring-shaped area around the centroid of each ROI (between 4-8px radius) and averaging the green fluorescence from these areas, obtaining a single-cell read-out of EGFP expression. To account for recording noise from session to session, this measure was normalized to the mean nuclear fluorescence of the H2BmCherry (see Supp. Fig. 10A). For the analyses of the signal stability after sham/5-Ph-IAA treatment, we only included neurons with a least distance of 3px and a minimal initial normalized green fluorescence ≥0.3.

For the experiments of Fig. 4, for each image (FOV) we used a custom-written MATLAB software to manually identify representative circular ROIs on the PSD-95.FingR:EGFP and the PSD-95:HT:mAID+ JF635-HT images. Specifically, we separately estimated ROIs for neuropil areas of the image (showing no nuclei and a punctate PSD-95 signal) and background areas of the image (blood vessel areas with very low signal intensity; see Supp. Fig. 10B). For neuropil and background respectively, we computed the mean fluorescence from all ROIs of a given FOV in both channels, PSD-95.FingR:EGFP and PSD-95:HT:mAID+ JF635-HT. Lastly, we divided the mean neuropil fluorescence by the mean background fluorescence, yielding a normalized read-out of bulk synaptic PSD-95 signals (see Supp. Fig. 10B). We separately compared the bulk fluorescence values of the PSD-95.FingR:EGFP and the PSD-95:HT:mAID+ JF635-HT channel.

### Statistical analyses

Comparison between pairs of experimental groups of experiments carried out in cell culture were tested using two-tailed t-tests without assuming identical variances using Microsoft Excel.

Group differences between the auxin and sham treated animals in the in-vivo experiments were tested by applying two-sided Wilcoxon rank sum tests using the Matlab statistics toolbox software.

## Supporting information

Plasmid sequences

## Acknowledgements

We are grateful to Tamar Galateanu, Leonid Odesski and for their invaluable assistance and to all members of the Ziv, Rumpel and Brose labs for their support and assistance. The project was supported by grants from the Deutsche Forschungsgemeinschaft German-Israeli Project Cooperation (DIP) to SR (#RU 900/5-1), NB and NEZ (ZI 1039/1-1), the Israel Science Foundation (1470/18; 2322/23), the Rappaport Institute and the Allen and Jewel Prince Center for Neurodegenerative Disorders of the Brain to NEZ, and the Deutsche Forschungsgemeinschaft CRC1080-C05, Deutsche Forschungsgemeinschaft SPP 2041 Project #347573108 and Deutsche Forschungsgemeinschaft/Agence nationale de la recherche Project #431393205 to SR.

## Figure Legends

**Supplemental Figure 1.**
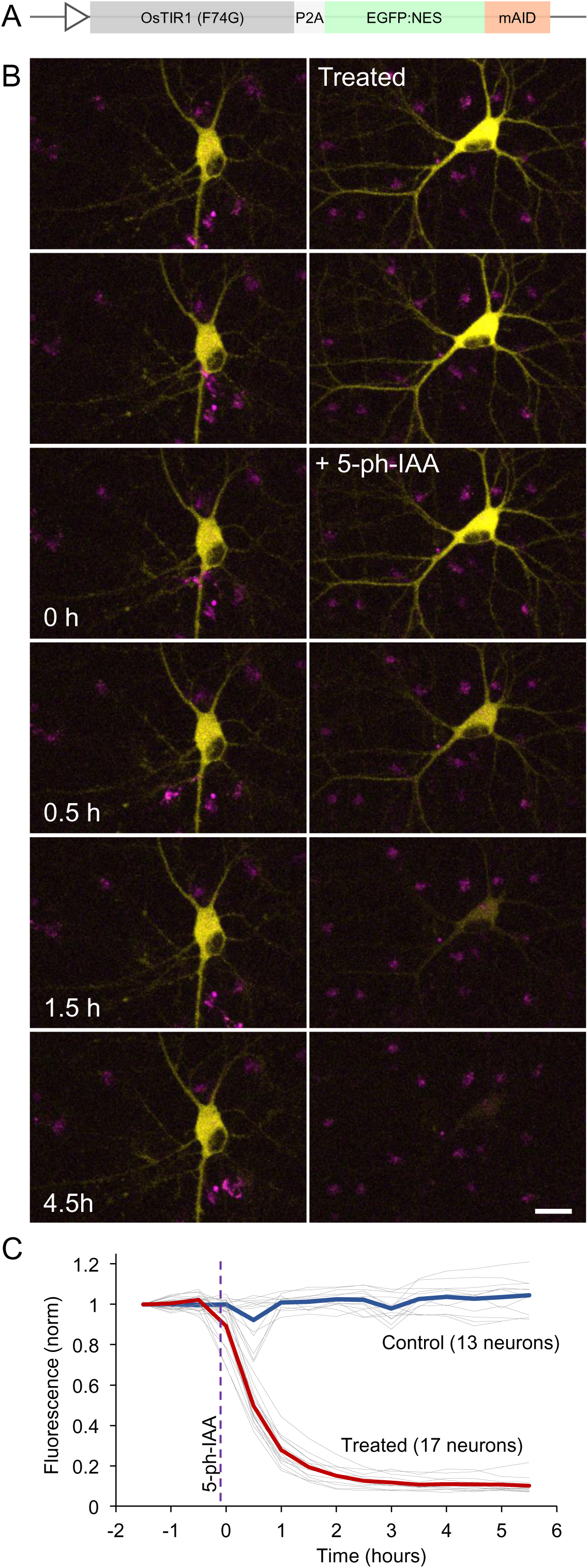
Rapid degradation of EGFP fused to a mAID degron. **A)** A vector for expressing both OsTIR(F74G) and EGFP fused to a mAID degron and a nuclear export sequence (NES), separated by a P2A sequence. Same construct as described by Yesbolatova et al., 2020, now in a lentiviral expression vector. **B)** Examples of neurons expressing OsTIR1(F74G) and EGFP:mAID (yellow) either exposed (right hand panels) or not exposed (left panels) to 200nM 5-Ph-IAA. Magenta dots are fluorescent background objects, used here to show that the loss of EGFP fluorescence is not a result of focal drift. Bar, 20µm. **C)** EGFP fluorescence measured at the cell bodies of 17 and 13 neurons exposed or not exposed to 5-Ph-IAA, respectively (thin gray lines). Fluorescence values for each neuron normalized to fluorescence measured at the last time point before 5-Ph-IAA was added. Thick red (5-Ph-IAA treated) and blue (untreated) lines represent population averages for each condition. Data from two separate experiments.

**Supplemental Figure 2.**
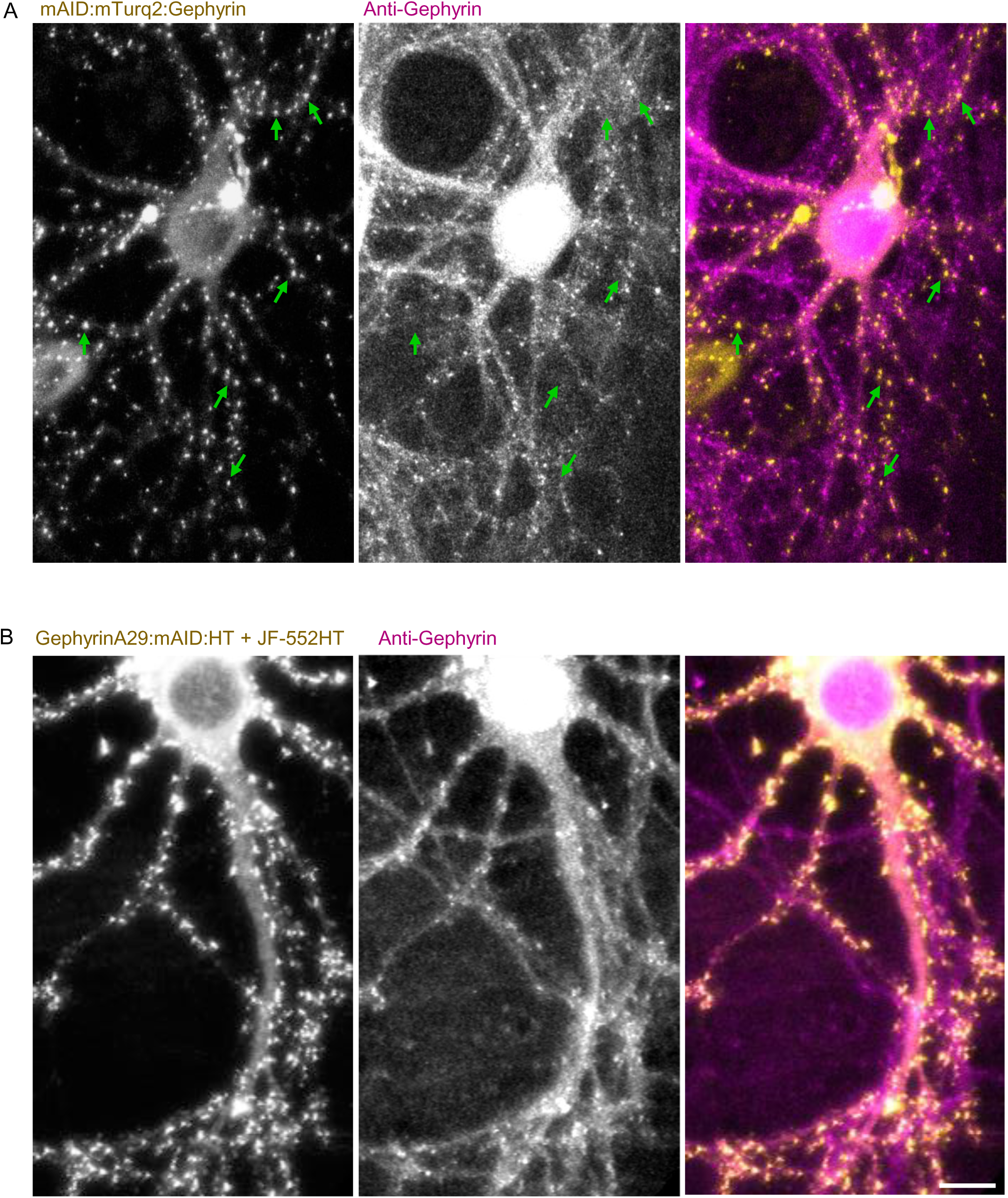
Recognition of gephyrin fusion proteins by anti-gephyrin antibodies. Neurons expressing mAID:mTurq2:Gephyrin or GephyrinA29:mAID:HT were fixed and labeled against gephyrin using an anti-gephyrin that recognizes the brain specific 93 kDa splice variant of gephyrin phosphorylated at Ser-270 (Synaptic Systems **#** 147 011). **A)** An example of a neuron expressing mAID:mTurq2:Gephyrin. Note that many mAID:mTurq2:Gephyrin puncta were not recognized by this antibody (some examples are shown by green arrows). **B)** An example of a neuron expressing GephyrinA29:mAID:HT labeled with the HaloTag ligand JF552-HT. Correspondence with antibody labeling is nearly perfect. Bar, 10 µm.

**Supplemental Figure 3.**
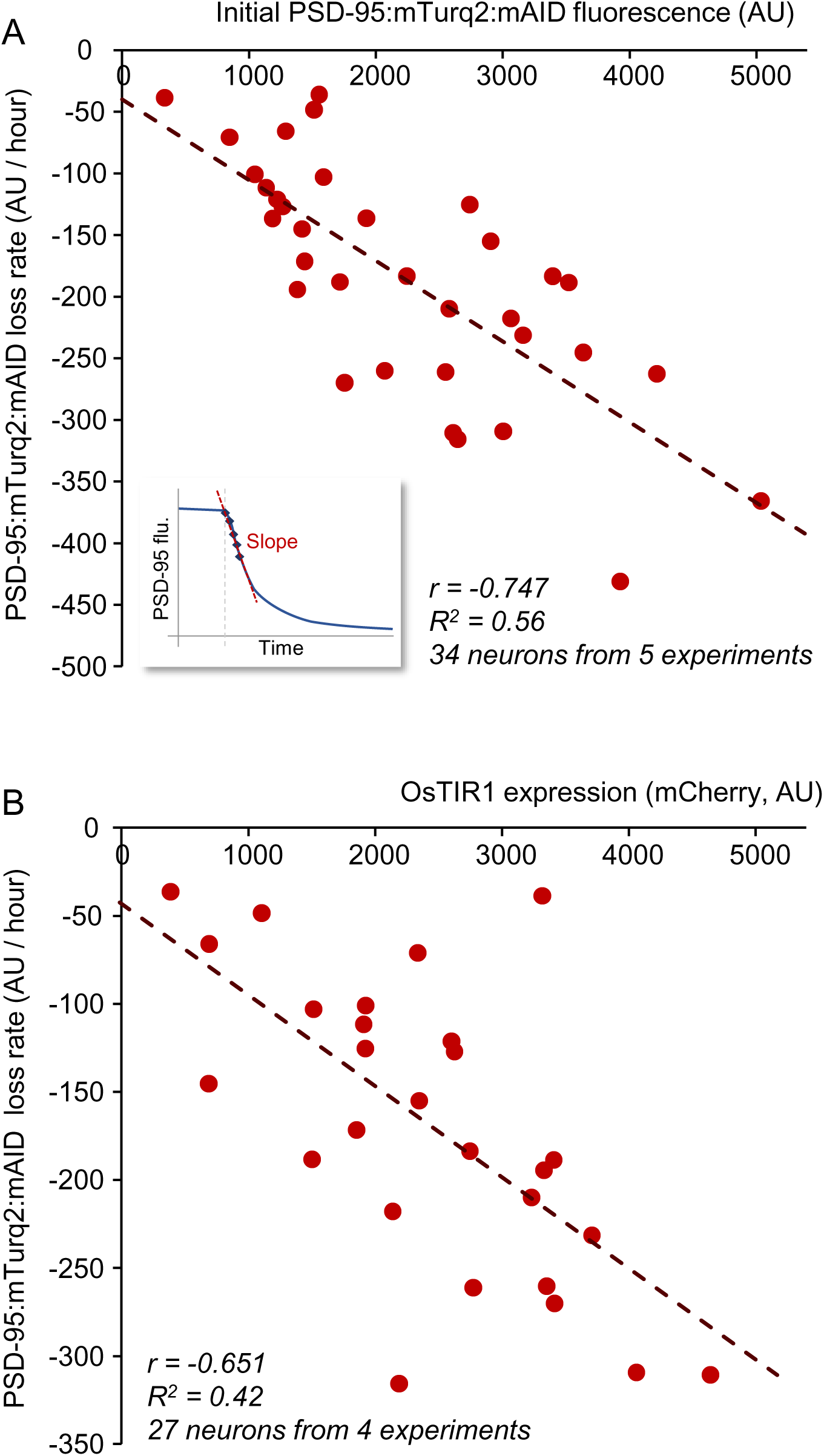
PSD-95:mTurq2:mAID loss rate dependencies. **A)** Rates of synaptic PSD-95:mTurq2:mAID loss (in fluorescence units / h) were calculated for each neuron coexpressing PSD-95:mTurq2:mAID and OsTIR1-P2A-mCherry by a fitting a line to mTurq2 fluorescence measurements made during the first 5 time points following exposure to 5-Ph-IAA (Inset). Loss rates were then plotted against initial PSD-95:mTurq2:mAID levels in the same neurons. The correlation between these two measures was -0.747 (34 neurons from 5 experiments). **B)** Similar plots comparing mTurq2 loss rate to mCherry fluorescence measured for the same neuron, with mCherry fluorescence serving as a surrogate for OsTIR1 expression. The correlation between these two measures was -0.651 (27 neurons from 4 experiments).

**Supplemental Figure 4.**
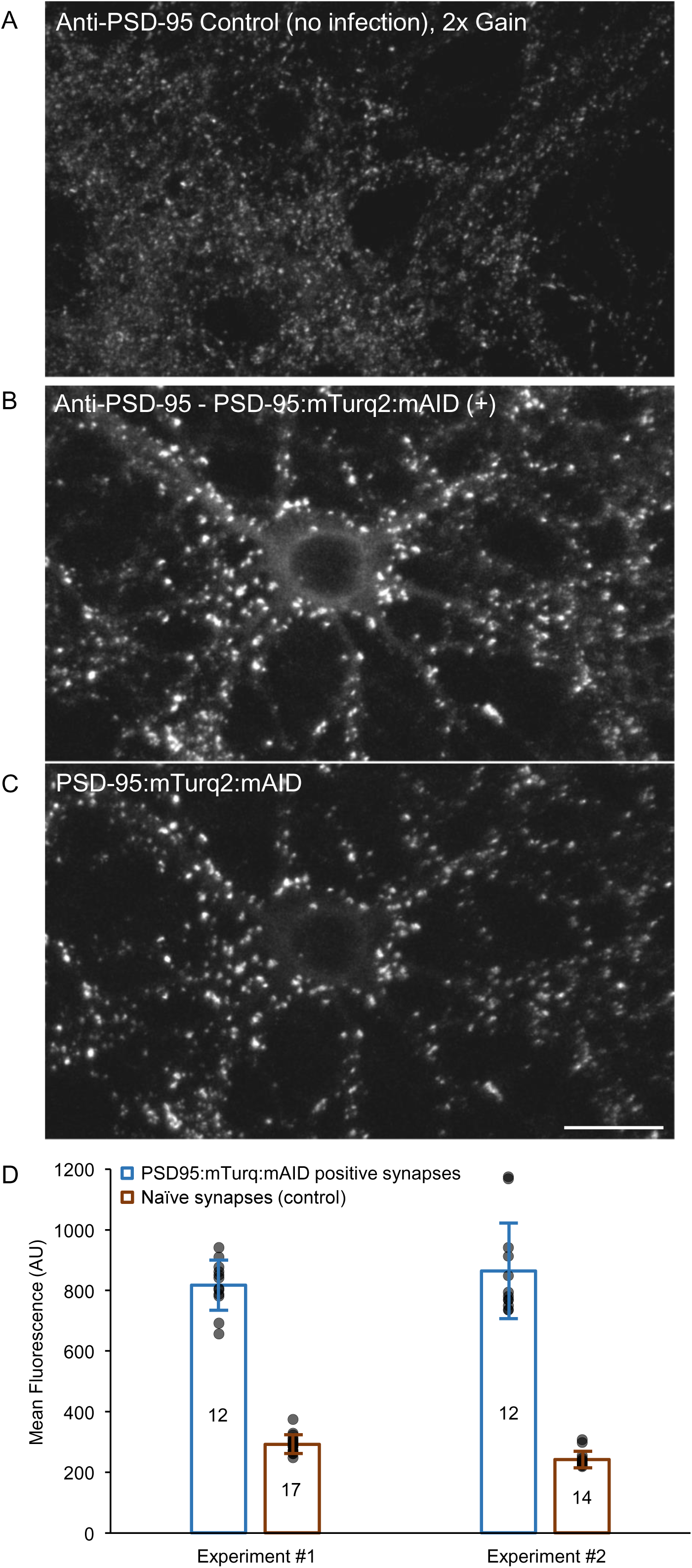
Quantification of PSD-95 overexpression. Networks of neurons expressing PSD-95:mTurq2:mAID, SEpH:GluA2 and OsTIR1-P2A-mCherry as well as age matched naïve networks from the same cell culture preparations were fixed and stained against PSD-95 (FluoTag®-X2 anti-PSD95 Alexa 647; NanoTag Biotechnologies #N3702-AF647-L; 1:200). **A)** Naïve (uninfected neurons) fixed and immunolabeled against PSD-95. **B)** A neuron expressing PSD-95:mTurq2:mAID fixed and immunolabeled against PSD-95. **C)** PSD-95:mTurq2:mAID fluorescence of the same synapses as in B. Note that in this particular field of view, all synapses were PSD-95:mTurq2:mAID positive. **D)** Comparison of anti-PSD-95 immunofluorescence of postsynaptic sites positive for PSD95:mTurq:mAID in the triple infected preparations to that of postsynaptic densities in the naïve ones (15,839 and 53,913 synapses from 24 and 31 fields of view, respectively; two separate experiments, shown separately). Every data point is the average fluorescence (arbitrary units) for one field of view. Columns and error bars show means and standard deviations. Correlations (Pearson’s) between PSD-95:mTurq2:mAID and anti-PSD-95 fluorescence on a synapse by synapses basis were 0.86 and 0.80 for the two experiments, indicating that anti-PSD-95 fluorescence provided a good readout of PSD-95 expression levels.

**Supplemental Figure 5.**
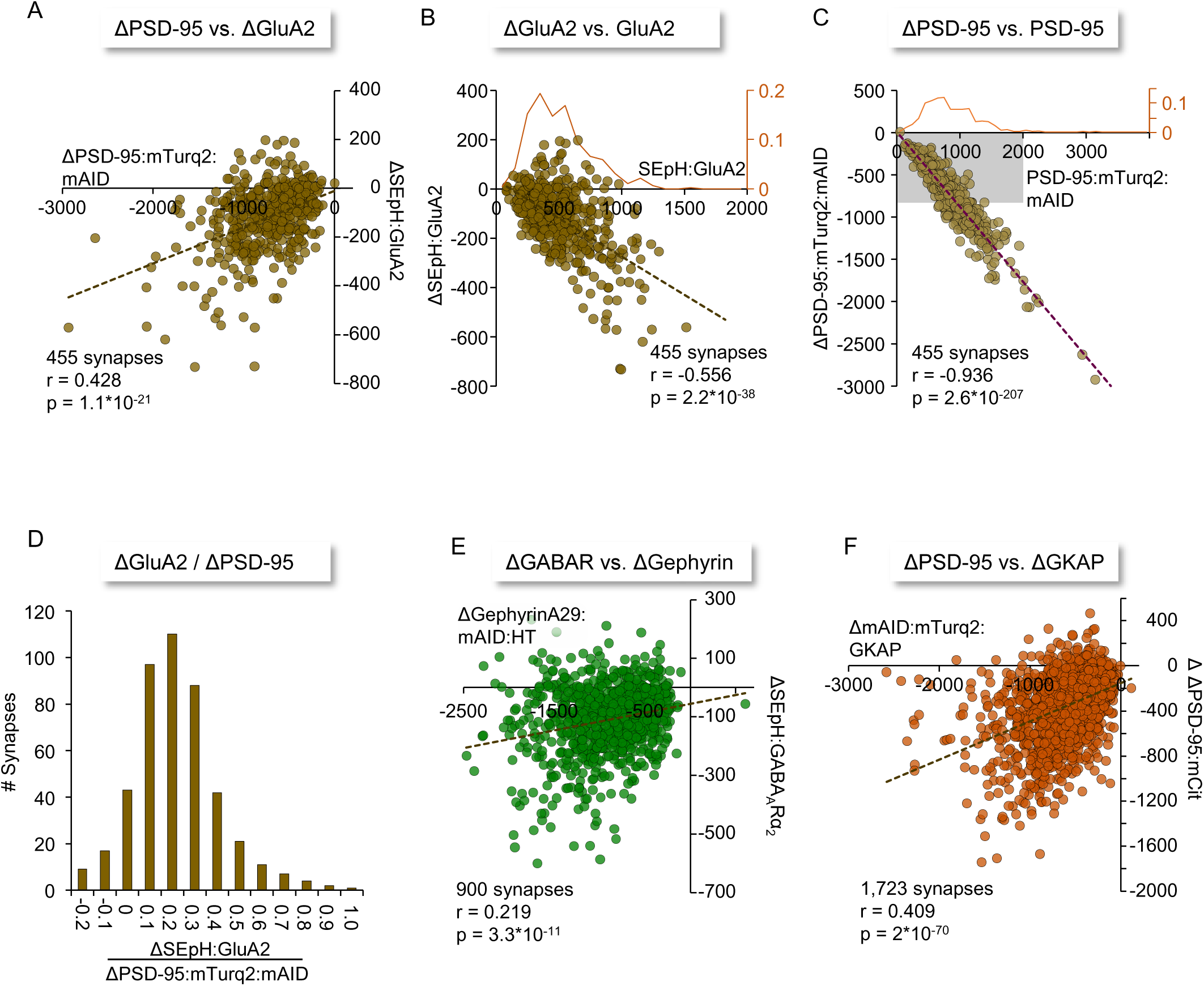
Change in protein contents of individual synapses. **A)** Comparisons, on a synapse by synapse basis, of changes in SEpH:GluA2 fluorescence as a function of PSD-95:mTurq2:mAID fluorescence loss at the same synapses measured 15 hours after exposure to 5-Ph-IAA. 455 synapses from 18 neurons in 3 separate experiments. Dashed line is a linear regression for these data, with its equation shown below. r = Pearson’s correlation. **B)** Dependence of changes in SEpH:GluA2 fluorescence as a function of initial SEpH:GluA2 fluorescence at the same synapses. Dashed line is a linear regression for these data, with its equation shown below. The scatter around the line, quantified by the coefficient of determination (R^2^) is 0.309. Orange line and right hand Y axis show the distribution of initial SEpH:GluA2 fluorescence values for these 455 synapses. **C)** Dependence of changes in PSD-95:mTurq2:mAID fluorescence as a function of initial PSD-95:mTurq2:mAID fluorescence at the same synapses. Dashed line is a linear regression for these data, with its equation shown below. Orange line and right hand Y axis shows the distribution of initial PSD-95:mTurq2:mAID fluorescence values for these 455 synapses. Note the absence of any synapse for which PSD-95:mTurq2:mAID fluorescence increased following 5-Ph-IAA exposure. Also note the tighter correlation and reduced scatter (R^2^ = 0.876) for these data as compared to B. Limiting the regression analysis to synapses with fluorescence levels comparable to those of SEpH:GluA2 in panel B (PSD-95:mTurq2:mAID no brighter than 2,000 fluorescence units and Δ fluorescence no greater that 800 units; gray rectangle; 271 synapses) resulted in a correlation of r = 0.826 (R^2^ = 0.682). The reduced scatter and tighter correlation for fluorescent objects of similar fluorescence levels indicates that the moderate dependence of SEpH:GluA2 loss on PSD-95:mTurq2:mAID (panel A), including observations of increased SEpH:GluA2 fluorescence, cannot be solely attributed to measurement noise. **D)** Distribution of ratios of SEpH:GluA2 fluorescence loss to PSD-95:mTurq2:mAID fluorescence loss at individual synapses. Negative values reflect gains of SEpH:GluA2 fluorescence. **E)** Comparisons, on a synapse by synapse basis, of changes in SEpH:GABA_A_Rα_2_ fluorescence as a function of GephyrinA29:mAID:HT fluorescence loss at the same synapses. 900 synapses from 18 neurons in 3 separate experiments. Dashed line is a linear regression for these data, with its equation is shown below. **F)** Comparisons, on a synapse by synapse basis, of changes in PSD-95:mCit fluorescence as a function of mAID:mTurq2:GKAP fluorescence loss at the same synapses 15 hours after exposure to 5-Ph-IAA. 1,723 synapses from 39 neurons in 5 separate experiments. Dashed line is a linear regression for these data, with its equation is shown below. All fluorescence data in this figure reflect averages of measurements made at three consecutive time points to minimize effects of measurement noise.

**Supplemental Figure 6.**
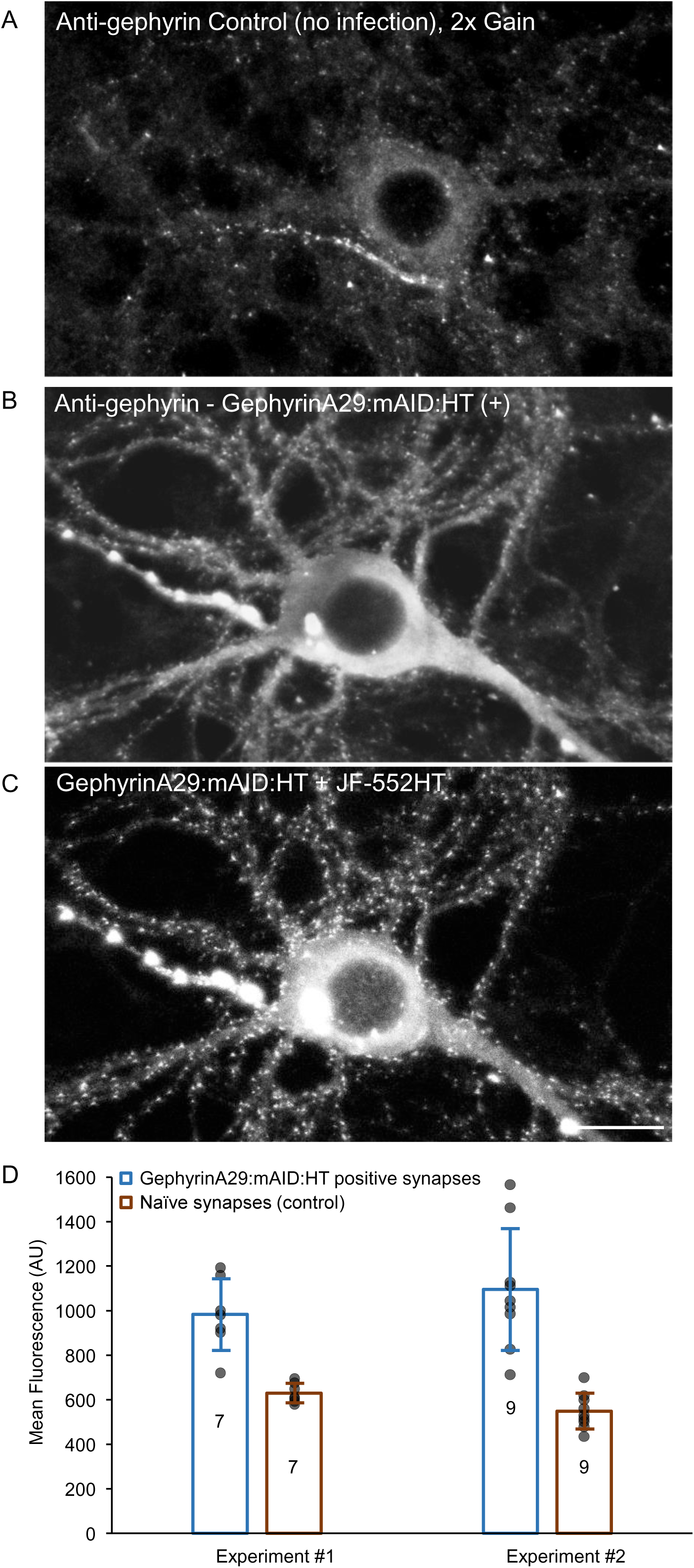
Quantification of Gephyrin overexpression. Preparations containing neurons expressing GephyrinA29:mAID:HT, SEpH:GABA_A_Rα_2_ and OsTIR1-P2A-mCherry as well as age matched naïve networks from the same cell culture preparations were fixed and stained against Gephyrin (monoclonal mouse anti gephyrin Synaptic Systems #147111). Infected networks were labeled before fixation with JF552-HT. **A)** Naïve (uninfected neurons) fixed and immunolabeled against gephyrin. **B)** A neuron expressing GephyrinA29:mAID:HT fixed and immunolabeled against gephyrin. **C)** GephyrinA29:mAID:HT + JF552-HT fluorescence of the same synapses as in B. **D)** Comparison of Anti-Gephyrin immunofluorescence of postsynaptic sites positive to JF552-HT fluorescence in the triple infected networks to that of postsynaptic densities in the naïve networks (6,204 and 4,626 synapses from 16 and 16 fields of view, respectively; two separate experiments, shown separately). Every data point is the average fluorescence (arbitrary units) for one field of view. Columns and error bars show means and standard deviations. Correlation (Pearson’s) between JF552-HT fluorescence and anti-Gephyrin fluorescence on a synapse by synapses basis was 0.50, indicating that anti-Gephyrin fluorescence provided a reasonable, if imperfect, readout of gephyrin expression levels, possibly affected by the use of a HT - HT ligand pair rather than a fluorescent protein

**Supplemental Figure 7.**
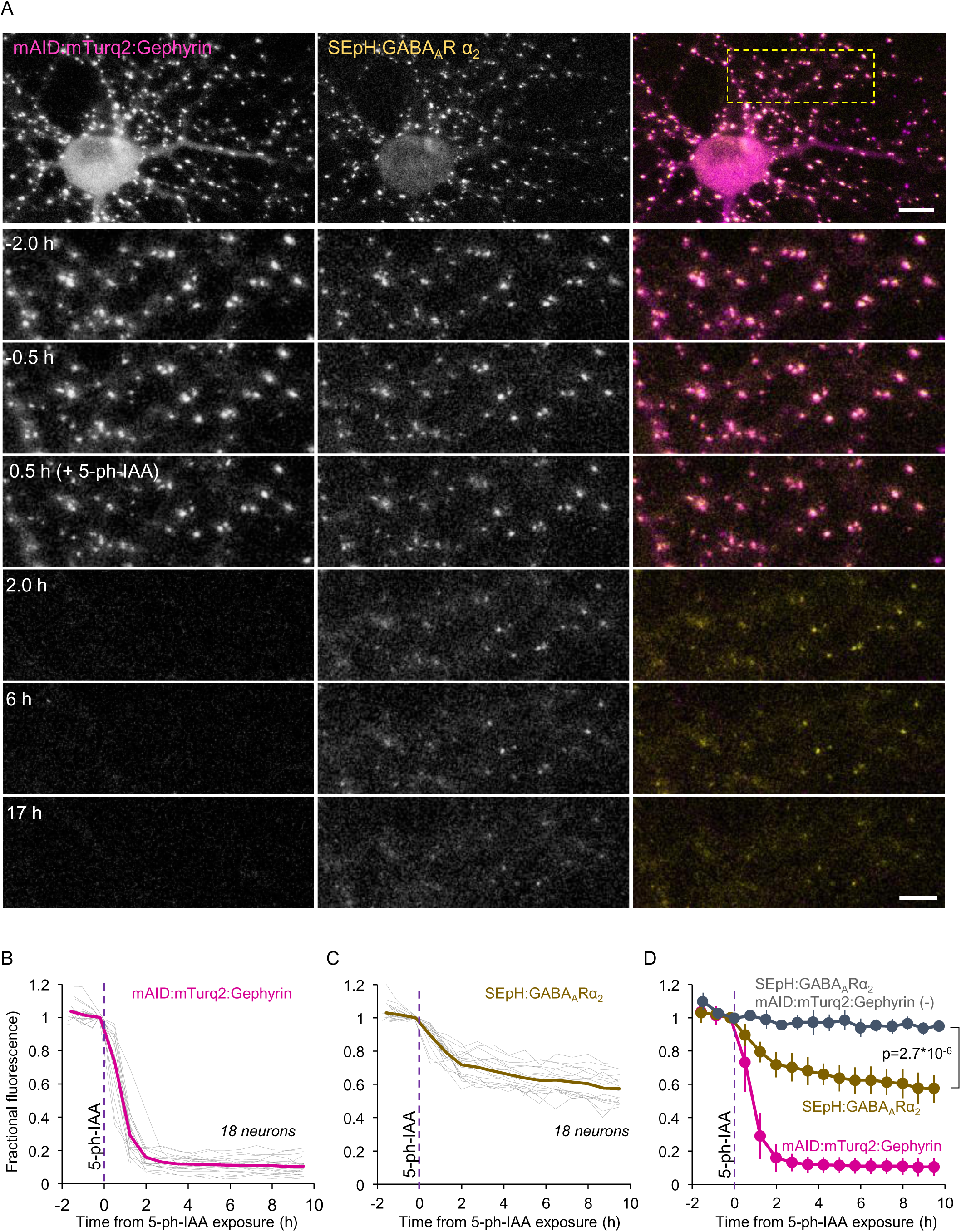
Acute elimination of mAID:mTurq2:Gephyrin is followed by loss of GABA receptors at the same synapses. **A)** Top panels: A rat cortical neuron in culture co-expressing mAID:mTurq2:Gephyrin, SEpH:GABA_A_Rα_2_ and OsTIR1-P2A-mCherry (not shown). Bottom panels: Region in yellow rectangle at greater detail, before, and after addition of 5-Ph-IAA. Note that the near complete loss of fluorescence (presumably reflecting mAID:mTurq2:Gephyrin degradation) is associated with a partial reduction in SEpH:GABA_A_Rα_2_ fluorescence. Scale bars: 10µm (top panels) 5µm (bottom panels). **B)** mAID:mTurq2:Gephyrin fluorescence measured at 50 synapses of each neuron tracked throughout the experiments. Each thin gray line is the average mAID:mTurq2:Gephyrin fluorescence (normalized to the time point just before 5-Ph-IAA addition) measured for the synapses of one neuron (18 neurons from 3 experiments). Thick magenta line is the population average**. C)** changes in SEpH:GABA_A_Rα_2_ fluorescence at the same synapses and neurons of B. Thick brown line is the population average. **D)** SEpH:GABA_A_Rα_2_ fluorescence measured at 200 synapses of neurons positive for SEpH:GABA_A_Rα_2_ and OsTIR1-P2A-mCherry but negative for mAID:mTurq2:Gephyrin. Each thin gray line is the average fluorescence measured for the synapses of one neuron (4 neurons from 1 experiment). Thick gray line is the population average. **E)** Pooled data. Error bars are standard deviations. Test for difference between mAID:mTurq2:Gephyrin positive and negative cells – unpaired t-test, without assuming equal variances; applied to data from last time point.

**Supplemental Figure 8.**
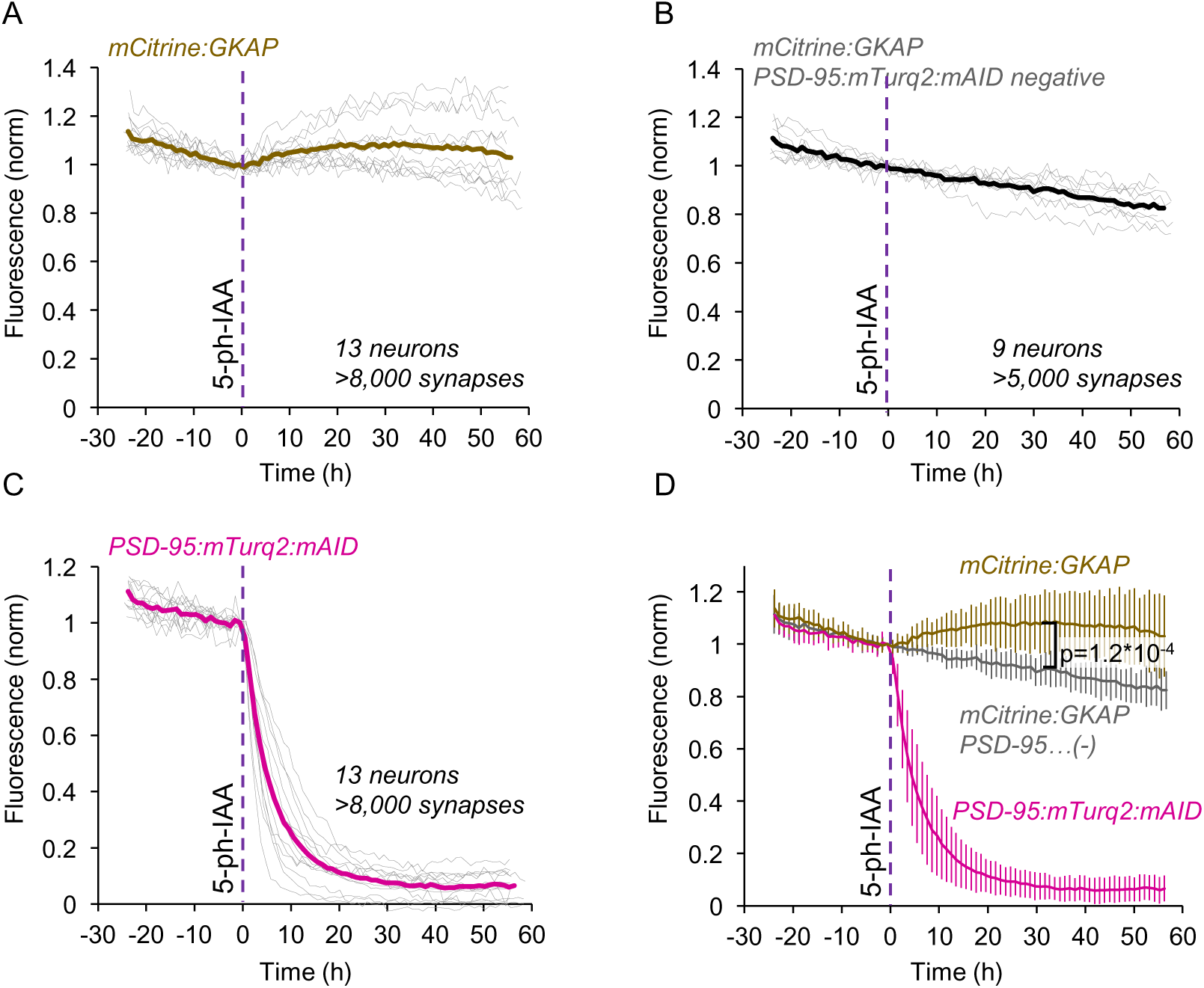
Acute elimination of PSD-95:mTurq2:mAID is followed by ‘substitution’ with GKAP. **A)** Cortical neurons in culture were infected with lentiviral expression vectors encoding for PSD-95:mTurq2:mAID, mCit:GKAP and OsTIR1-P2A-mCherry. Changes in mCit:GKAP fluorescence were measured at synapses in fields of view containing neurons expressing all three exogenous proteins. Here, unlike Fig. 7, synapses were not tracked individually but located programmatically anew at each time step, resulting in an unbiased selection of synapses but with no distinction between synapses belonging to triple expressing neurons and synapses belonging to other neurons in the same fields of view. Each thin gray line is the average fluorescence measured for the synapses in one field of view (13 fields of view from 3 experiments). Thick brown line is the population average. **B)** mCit:GKAP fluorescence measured at synapses in fields of view containing neurons positive for mCit:GKAP and OsTIR1-P2A-mCherry but negative for PSD-95:mTurq2:mAID. Each thin gray line is the average fluorescence measured for the synapses of one neuron (9 neurons from 3 experiments). Thick gray line is the population average. **C)** PSD-95:mTurq2:mAID fluorescence measured at the same synapses and neurons of A. Thick magenta line is the population average. **D)** Pooled data. Error bars are standard deviations. Test for difference between PSD-95:mTurq2:mAID positive and negative cells – unpaired t-test, without assuming equal variances; applied to data from t=30 h.

**Supplemental Figure 9.**
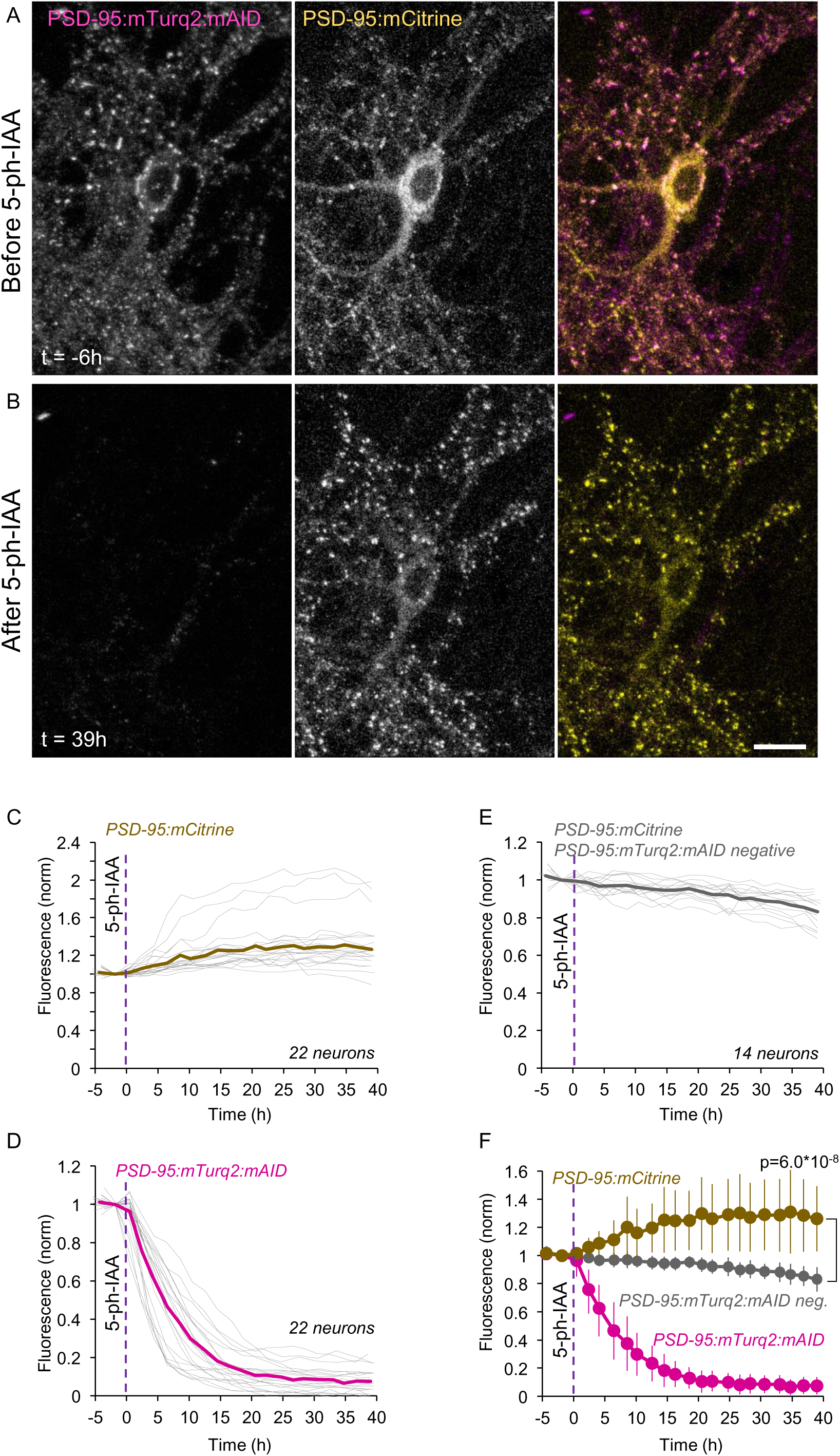
Acute elimination of PSD-95:mTurq2:mAID is followed by rapid replacement with PSD-95:mCitrine. **A)** A rat cortical neuron in culture co-expressing PSD-95:mTurq2:mAID (left) and PSD-95:mCitrine (middle) as well as OsTIR1-P2A-mCherry (not shown). **B)** 5-Ph-IAA induced nearly complete loss of PSD-95:mTurq2:mAID, and increased synaptic levels of PSD-95:mCitrine at the same synapses. Scale bar: 20 µm. **C)** Changes in PSD-95:mCitrine fluorescence measured at 15-34 synapses of each neuron tracked throughout the experiments (485 in total). Each thin gray line is the average normalized fluorescence measured for the synapses of one neuron (22 neurons from 3 experiments). Thick brown line is the population average. **D)** PSD-95:mTurq2:mAID fluorescence measured at the same synapses and neurons of C. Thick magenta line is the population average. **E)** PSD-95:mCitrine fluorescence measured at 14-42 synapses of neurons positive for PSD-95:mCitrine and OsTIR1-P2A-mCherry but negative for PSD-95:mTurq2:mAID (380 in total). Each thin gray line is the average fluorescence measured for the synapses of one neuron (14 neurons from 3 experiments). Thick gray line is the population average. **F)** Pooled data. Error bars are standard deviations. Test for difference between PSD-95:mTurq2:mAID positive and negative cells – unpaired t-test, without assuming equal variances; applied to data from last time point.

**Supplemental Figure 10.**
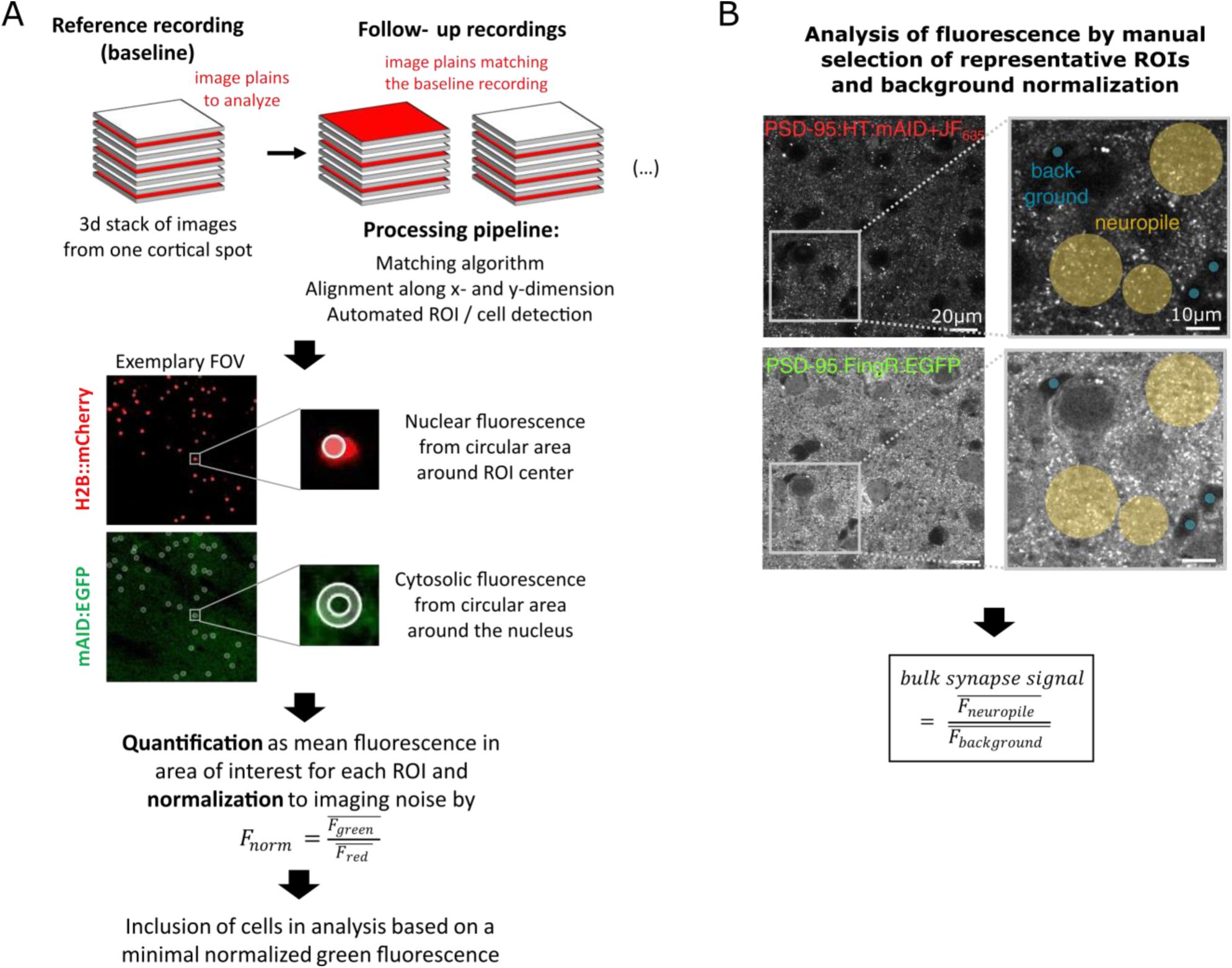
Data processing and analysis of *in-vivo* experiments: **A)** Processing and analysis pipeline for two-photon image stacks to quantify expression of the cytosolically expressed mAID:EGFP (Fig. 3). Single image planes (syn. fields of view, FOV) were matched and aligned across all imaging time points. Cytosolic regions of interest (ROIs) were identified as ring-shaped areas round the nuclear H2B:mCherry blobs. The single-cell EGFP signal was quantified normalizing the mean green cytosolic fluorescence by the mean nuclear red fluorescence. Only cells with a reliable baseline signal (F_norm_ ≥ 0.3) were included in the analysis (see Methods). **B)** Analysis of synaptic PSD-95 signals (Fig. 4). For both the PSD-95:FingR:EGFP signal and the PSD-95:JF635-HT signal, neuropil fluorescence was quantified by manually selecting representative circular ROIs from the images (yellow circles). Moreover, background signals were assessed by estimating fluorescence in ROIs on blood vessels (blue circles). The bulk synaptic PSD-95 signal was then quantified as the mean background-normalized neuropil fluorescence.

## References

Ashby MC, De La Rue SA, Ralph GS, Uney J, Collingridge GL, Henley JM. (2004) Removal of AMPA receptors (AMPARs) from synapses is preceded by transient endocytosis of extrasynaptic AMPARs. J Neurosci. 24:5172–5176.

Bai G, Wang Y, Zhang M. (2021) Gephyrin-mediated formation of inhibitory postsynaptic density sheet via phase separation. Cell Res. 31:312–325.

Bai G, Zhang M. Inhibitory postsynaptic density from the lens of phase separation. (2022) Oxf Open Neurosci. 1:kvac003.

Béïque JC, Andrade R. (2003) PSD-95 regulates synaptic transmission and plasticity in rat cerebral cortex. J Physiol. 546(Pt 3):859–867.

Béïque JC, Lin DT, Kang MG, Aizawa H, Takamiya K, Huganir RL. (2006) Synapse-specific regulation of AMPA receptor function by PSD-95. Proc Natl Acad Sci U S A. 103:19535–19540.

Bessa-Neto D, Choquet D. (2023) Molecular mechanisms of AMPAR reversible stabilization at synapses. Mol Cell Neurosci. 125:103856.

Bondeson DP, Mullin-Bernstein Z, Oliver S, Skipper TA, Atack TC, Bick N, Ching M, Guirguis AA, Kwon J, Langan C, Millson D, Paolella BR, Tran K, Wie SJ, Vazquez F, Tothova Z, Golub TR, Sellers WR, Ianari A. (2022) Systematic profiling of conditional degron tag technologies for target validation studies. Nat Commun. 13:5495.

Bulovaite E, Qiu Z, Kratschke M, Zgraj A, Fricker DG, Tuck EJ, Gokhale R, Koniaris B, Jami SA, Merino-Serrais P, Husi E, Mendive-Tapia L, Vendrell M, O’Dell TJ, DeFelipe J, Komiyama NH, Holtmaat A, Fransén E, Grant SGN. (2022) A brain atlas of synapse protein lifetime across the mouse lifespan. Neuron 110:4057–4073.

Calamai M, Specht CG, Heller J, Alcor D, Machado P, Vannier C, Triller A. (2009) Gephyrin oligomerization controls GlyR mobility and synaptic clustering. J Neurosci. 29:7639–7648.

Chen X, Nelson CD, Li X, Winters CA, Azzam R, Sousa AA, Leapman RD, Gainer H, Sheng M, Reese TS. (2011) PSD-95 is required to sustain the molecular organization of the postsynaptic density. J Neurosci. 31:6329–6338.

Chen X, Wu X, Wu H, Zhang M. (2020) Phase separation at the synapse. Nat Neurosci. 23:301-310. Choii G, Ko J. (2015) Gephyrin: a central GABAergic synapse organizer. Exp Mol Med. 47:e158.

Cohen LD, Ziv NE. Neuronal and synaptic protein lifetimes (2019) Curr Opin Neurobiol. 57:9–16.

Dvorkin R, Ziv NE. (2016) Relative Contributions of Specific Activity Histories and Spontaneous Processes to Size Remodeling of Glutamatergic Synapses. PLoS Biol. 14:e1002572.

Ehrlich I, Klein M, Rumpel S, Malinow R. (2007) PSD-95 is required for activity-driven synapse stabilization. Proc Natl Acad Sci U S A. 104:4176–4181.

Ehrlich I, Malinow R. (2004) Postsynaptic density 95 controls AMPA receptor incorporation during long-term potentiation and experience-driven synaptic plasticity. J Neurosci. 24:916–927.

El-Husseini AE, Schnell E, Chetkovich DM, Nicoll RA, Bredt DS. (2000) PSD-95 involvement in maturation of excitatory synapses. Science 290:1364–1368.

Elias GM, Funke L, Stein V, Grant SG, Bredt DS, Nicoll RA (2006). Synapse-specific and developmentally regulated targeting of AMPA receptors by a family of MAGUK scaffolding proteins. Neuron. 52:307–320.

England CG, Luo H, Cai W. (2015) HaloTag technology: a versatile platform for biomedical applications. Bioconjug Chem. 26:975–986.

Eppler, JB (2022) 2-photon data preprocessing pipeline. DOI 10.5281/zenodo.5822485.

Fan X, Jin WY, Lu J, Wang J, Wang YT (2014). Rapid and reversible knockdown of endogenous proteins by peptide-directed lysosomal degradation. Nat Neurosci. 17:471–480.

Futai K, Kim MJ, Hashikawa T, Scheiffele P, Sheng M, Hayashi Y. (2007) Retrograde modulation of presynaptic release probability through signaling mediated by PSD-95-neuroligin. Nat Neurosci. 10:186–195.

Goedhart J, van Weeren L, Hink MA, Vischer NO, Jalink K, Gadella TW Jr. (2010) Bright cyan fluorescent protein variants identified by fluorescence lifetime screening. Nat Methods. 7:137–139.

Goel K, Ploski JE. (2022) RISC-y Business: Limitations of Short Hairpin RNA-Mediated Gene Silencing in the Brain and a Discussion of CRISPR/Cas-Based Alternatives. Front Mol Neurosci. 15:914430.

Grimm JB, Muthusamy AK, Liang Y, Brown TA, Lemon WC, Patel R, Lu R, Macklin JJ, Keller PJ, Ji N, Lavis LD. (2017) A general method to fine-tune fluorophores for live-cell and in vivo imaging. Nat Methods 14:987–994.

Gross GG, Junge JA, Mora RJ, Kwon HB, Olson CA, Takahashi TT, Liman ER, Ellis-Davies GC, McGee AW, Sabatini BL, Roberts RW, Arnold DB. (2013) Recombinant probes for visualizing endogenous synaptic proteins in living neurons. Neuron 78:971–985.

Gross GG, Straub C, Perez-Sanchez J, Dempsey WP, Junge JA, Roberts RW, Trinh le A, Fraser SE, De Koninck Y, De Koninck P, Sabatini BL, Arnold DB. (2016) An E3-ligase-based method for ablating inhibitory synapses. Nat Methods. 13:673–678.

Hammond S, Caudy A, Hannon G. (2001) Post-transcriptional gene silencing by double-stranded RNA. Nat Rev Genet 2, 110–119.

Hayashi Y, Ford LK, Fioriti L, McGurk L, Zhang M. (2021) Liquid-Liquid Phase Separation in Physiology and Pathophysiology of the Nervous System. J Neurosci. 41:834–844.

Kim E, Naisbitt S, Hsueh YP, Rao A, Rothschild A, Craig AM, Sheng M. (1997) GKAP, a novel synaptic protein that interacts with the guanylate kinase-like domain of the PSD-95/SAP90 family of channel clustering molecules. J Cell Biol. 136:669–678.

Kim MJ, Futai K, Jo J, Hayashi Y, Cho K, Sheng M. (2007) Synaptic accumulation of PSD-95 and synaptic function regulated by phosphorylation of serine-295 of PSD-95. Neuron 56:488–502.

Kopec CD, Li B, Wei W, Boehm J, Malinow R (2006) Glutamate receptor exocytosis and spine enlargement during chemically induced long-term potentiation. J Neurosci 26:2000–2009.

Kuriu T, Inoue A, Bito H, Sobue K, Okabe S. (2006) Differential control of postsynaptic density scaffolds via actin-dependent and -independent mechanisms. J Neurosci. 26:7693–7706.

Levy JM, Chen X, Reese TS, Nicoll RA. (2015) Synaptic Consolidation Normalizes AMPAR Quantal Size following MAGUK Loss. Neuron 87:534–548.

Lois C, Hong EJ, Pease S, Brown EJ, Baltimore D. (2002) Germline transmission and tissue-specific expression of transgenes delivered by lentiviral vectors. Science. 295:868–872.

Machado P, Rostaing P, Guigonis JM, Renner M, Dumoulin A, Samson M, Vannier C, Triller A. (2011) Heat shock cognate protein 70 regulates gephyrin clustering. J Neurosci. 31:3–14.

Makino-Itou H, Yamatani N, Okubo A, Kiso M, Ajima R, Kanemaki MT, Saga Y. (2024) Establishment and characterization of mouse lines useful for endogenous protein degradation via an improved auxin-inducible degron system (AID2). Dev Growth Differ. 66:384–393.

Migaud M, Charlesworth P, Dempster M, Webster LC, Watabe AM, Makhinson M, He Y, Ramsay MF, Morris RG, Morrison JH, O’Dell TJ, Grant SG. (1998) Enhanced long-term potentiation and impaired learning in mice with mutant postsynaptic density-95 protein. Nature. 396:433–439.

Mohar B, Grimm JB, Patel R, Brown TA, Tillberg PW, Lavis LD, Spruston N, Svoboda K (2023) Brain-wide measurement of protein turnover with high spatial and temporal resolution. bioRxiv 2022.11.12.516226; doi: 10.1101/2022.11.12.516226

Muller, S (2023) Characterization of a c-Fos reporter system for in vivo imaging in the mouse auditory cortex (Thesis). 10.25358/openscience-9595

Nakamura Y, Morrow DH, Modgil A, Huyghe D, Deeb TZ, Lumb MJ, Davies PA, Moss SJ. (2016) Proteomic Characterization of Inhibitory Synapses Using a Novel pHluorin-tagged γ-Aminobutyric Acid Receptor, Type A (GABAA), α2 Subunit Knock-in Mouse. J Biol Chem. 291:12394–12407.

Nakano R, Ihara N, Morikawa S, Nakashima A, Kanemaki MT, Ikegaya Y, Takeuchi H. (2019) Auxin-mediated rapid degradation of target proteins in hippocampal neurons. Neuroreport. 30:908–913.

Natsume T, Kanemaki MT. (2017) Conditional Degrons for Controlling Protein Expression at the Protein Level. Annu Rev Genet. 51:83–102.

Nishimura K, Fukagawa T, Takisawa H, Kakimoto T, Kanemaki M. (2009) An auxin-based degron system for the rapid depletion of proteins in nonplant cells. Nat Methods 6:917–922.

Rasmussen AH, Rasmussen HB, Silahtaroglu A. (2017) The DLGAP family: neuronal expression, function and role in brain disorders. Mol Brain. 10:43.

Rubinski A, Ziv NE. (2015) Remodeling and Tenacity of Inhibitory Synapses: Relationships with Network Activity and Neighboring Excitatory Synapses. PLoS Comput Biol. 11:e1004632.

Salvatico C, Specht CG, Triller A. (2015) Synaptic receptor dynamics: from theoretical concepts to deep quantification and chemistry *in cellulo*. Neuropharmacology 88:2–9.

Schlüter OM, Xu W, Malenka RC. (2006) Alternative N-terminal domains of PSD-95 and SAP97 govern activity-dependent regulation of synaptic AMPA receptor function. Neuron. 51:99–111.

Shi H, Jin Q, Chen F, Ouyang Z, Gou S, Liu X, Li L, Mu S, Lai C, Zhang Q, Ye Y, Wang K, Lai L (2022) CIRKO: A chemical-induced reversible gene knockout system for studying gene function in situ. bioRxiv 2022.08.31.506064; doi: 10.1101/2022.08.31.506064

Shin SM, Zhang N, Hansen J, Gerges NZ, Pak DT, Sheng M, Lee SH. (2012) GKAP orchestrates activity-dependent postsynaptic protein remodeling and homeostatic scaling. Nat Neurosci. 15:1655–1666.

Tretter V, Jacob TC, Mukherjee J, Fritschy JM, Pangalos MN, Moss SJ. (2008) The clustering of GABA(A) receptor subtypes at inhibitory synapses is facilitated via the direct binding of receptor alpha 2 subunits to gephyrin. J Neurosci. 28 :1356–1365.

Tsai JM, Nowak RP, Ebert BL, Fischer ES. (2024) Targeted protein degradation: from mechanisms to clinic. Nat Rev Mol Cell Biol. Apr 29. doi: 10.1038/s41580-024-00729-9. Epub ahead of print.

Tsai JM, Nowak RP, Ebert BL, Fischer ES. (2024) Targeted protein degradation: from mechanisms to clinic. Nat Rev Mol Cell Biol. 25:740–757.

Vistrup-Parry M, Chen X, Johansen TL, Bach S, Buch-Larsen SC, Bartling CRO, Ma C, Clemmensen LS, Nielsen ML, Zhang M, Strømgaard K. (2021) Site-specific phosphorylation of PSD-95 dynamically regulates the postsynaptic density as observed by phase separation. iScience. 4(11):103268.

Won S, Levy JM, Nicoll RA, Roche KW. MAGUKs: multifaceted synaptic organizers. (2017) Curr Opin Neurobiol. 43:94–101.

Yao I, Iida J, Nishimura W, Hata Y. (2003) Synaptic localization of SAPAP1, a synaptic membrane-associated protein. Genes Cells. 8:121–129.

Yesbolatova A, Saito Y, Kitamoto N, Makino-Itou H, Ajima R, Nakano R, Nakaoka H, Fukui K, Gamo K, Tominari Y, Takeuchi H, Saga Y, Hayashi KI, Kanemaki MT. (2020) The auxin-inducible degron 2 technology provides sharp degradation control in yeast, mammalian cells, and mice. Nat Commun. 11:5701.

Yudowski GA, Olsen O, Adesnik H, Marek KW, Bredt DS. (2013) Acute inactivation of PSD-95 destabilizes AMPA receptors at hippocampal synapses. PLoS One 8:e53965.

Zeidan A, Ziv NE. (2012) Neuroligin-1 loss is associated with reduced tenacity of excitatory synapses. PLoS One. 7:e42314.

Zeng M, Chen X, Guan D, Xu J, Wu H, Tong P, Zhang M. (2018) Reconstituted Postsynaptic Density as a Molecular Platform for Understanding Synapse Formation and Plasticity. Cell. 174:1172–1187.

Zeng M, Díaz-Alonso J, Ye F, Chen X, Xu J, Ji Z, Nicoll RA, Zhang M. (2019) Phase Separation-Mediated TARP/MAGUK Complex Condensation and AMPA Receptor Synaptic Transmission. Neuron 104:529–543.e6.

Zhu F, Cizeron M, Qiu Z, Benavides-Piccione R, Kopanitsa MV, Skene NG, Koniaris B, DeFelipe J, Fransén E, Komiyama NH, Grant SGN. (2018) Architecture of the Mouse Brain Synaptome. Neuron 99:781–799.

Zhu J, Zhou Q, Shang Y, Li H, Peng M, Ke X, Weng Z, Zhang R, Huang X, Li SSC, Feng G, Lu Y, Zhang M. (2017) Synaptic Targeting and Function of SAPAPs Mediated by Phosphorylation-Dependent Binding to PSD-95 MAGUKs. Cell Rep. 21:3781–3793.

Zhu S, Shen Z, Wu X, Han W, Jia B, Lu W, Zhang M. (2024) Demixing is a default process for biological condensates formed via phase separation. Science. 384:920–928.

